# Bone mechano-response is driven by locomotion transitions during vertebrate evolution

**DOI:** 10.1101/2024.08.29.610233

**Authors:** Saeka Shimochi, Clara Brunet, Margalida Fontcuberta-Rigo, Katja Hrovat, Pere Puigbò, Miho Nakamura

## Abstract

The skeleton supports the muscles in keeping the body upright against gravity while enduring thousands of daily loads. In this study, we investigated non-collagenous bone matrix proteins using osteoblast cell cultures and phylogenetic analyses to identify the molecular mechanisms involved in mechanical loading. The results indicate that the bone mechano-response is an evolutionary-driven process and that several non-collagenous proteins may significantly regulate the bone’s response to mechanical stress. According to our results, two significant evolutionary transitions in vertebrate locomotion shaped the roles of non-collagenous proteins in humans: the water-to-land transition, which increased mechanical stress on the limbs, and the evolution to bipedalism in humans, which altered the distribution of stress on the lower and upper limbs. Fetuin A, positively selected in both evolutionary transitions, showed the most significant expression change during mechanical stimulation.

## Main Text

The skeleton’s main mechanical function is to provide support for muscles to keep the body upright against gravity (*1*). Furthermore, many bones are subjected to thousands of repetitive loads (impacts) each day (*1*). Mechanical loading is essential for bone health and development because it plays a critical role in maintaining bone structure, density, and strength (*2*). Alterations in the skeleton dynamic process of bone resorption and formation (*1*) lead to bone diseases, including osteoporosis, the most common bone problem in elderly populations. A reduction in mechanical loading due to prolonged bed rest (*3*) or long-term exposure to microgravity (*4*) can lead to a reduction in bone mass. This understanding supports the general recommendation of weight-bearing exercise (together with getting adequate calcium, sunlight, and vitamin D (*5*)) to decrease osteoporosis risk (*6*). However, the molecular mechanism through which weight-bearing exercises prevent osteoporosis are not fully understood. Several studies in the field of mechanobiology have determined the roles of some cell types, cellular membrane receptors, and bone extracellular matrix (ECM) proteins in mechanical loading (*1, 7–9*). Osteocytes function as mechanosensing cells in bone tissue by transduction of mechanical signals to biochemical responses (*7*). PIEZO1 as a mechanosensing protein (*10*) regulates homeostasis via crosstalk of osteoblasts and osteoclasts (*8*). Osteopontin is a non-collagenous protein in the ECM related to mechanical stress (*9*). However, bone ECM proteome comprises hundreds of proteins (*11*) that are poorly understood and may play a major regulatory role in bone responses to mechanical loading (table S1).

Wolff’s law postulates that bone tissue can convert mechanical loading into structural change (*12*). Previous research reported that electricity is generated in collagen fibrils, which represent 90% of the organic components of the bone ECM, during weight-bearing exercises (*13*). In a previous study, we showed that the electricity produced in collagen fibrils is stored in the inorganic components of the ECM (*14*). Since bone ECM plays both structural role as a mechanical support and regulatory role as a key component of stem cell niches, we hypothesize that non-collagenous organic components of the bone matrix should have a key function in bone regulation through mechanical loading. Moreover, we hypothesized that two significant locomotion transitions in the evolution of vertebrates shaped the role of no-ncollagenous proteins in humans. The first locomotion transition was the water-to-land transition, which occurred at the time of origin of the tetrapods (approximately 400 Ma), and involved stronger mechanical stress on the limbs of tetrapods, in addition to ecological, cognitive, and physiological adaptations (*15*). The second locomotion transition occurred during the evolution to bipedalism in the hominini (approximately 11.6 Ma) and was characterized by a striking skeletal adaptation for walking upright on two feet (*16, 17*), leading to a differential mechanical stress on arms (less stress) and legs (more stress).

### Testing the conservation hypothesis

Human (*Homo sapiens*) is the only vertebrate to have acquired the spinal function to support the body against gravity and achieve upright bipedal locomotion (*18*). Here, we hypothesize that poorly conserved bone ECM proteins, with a large number of adaptive mutations during major locomotion transitions in the evolution of vertebrates, play a major role in bone mechano-responses to mechanical loading in humans. Zebrafish (*Danio rerio*) and human have 251 (out of 255) bone ECM homologous proteins (*11*). These proteins show a wide range of amino acids conservation (percentage of identity in pairwise sequence alignments from 0.00% to 99.86%) (Fig. 1A, table S2). Highly conserved homologous proteins between zebrafish and human are also structurally conserved (table S3). Furthermore, some homologous proteins with poorly conserved amino acid sequences are structurally conserved due to functional constraints of the protein (e.g. Fetuin A) (Fig. 1AB, table S4).

**Fig. 1.**
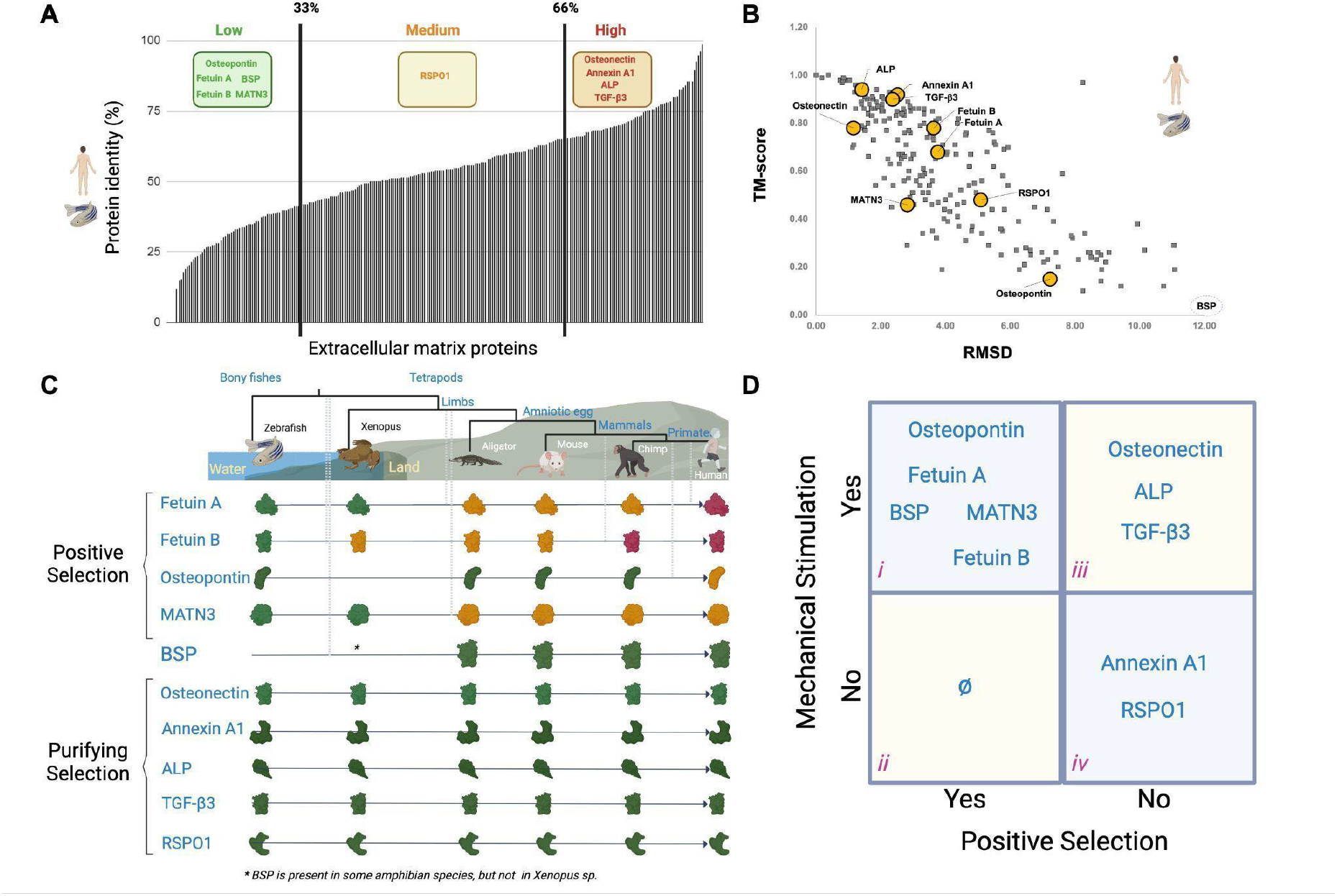
Analysis of the evolution of bone ECM proteins in vertebrates. **(A)** Percentage identity of bone ECM homologous proteins between *H. sapiens* and *D. rerio* (table S2). **(B)** Correlation of the Root Mean Square deviation (RMSD) and the template modeling score (TM-Score) of bone ECM homologous proteins between *Homo sapiens* and *Danio rerio* (full results are available in supplementary table S3). **(C)** Schematic representation of the positive selection analysis of bone ECM proteins in vertebrate evolution (full results are available in tables S4-6). **(D)** Contingency table of mechanical stimulation and positive selection. The table is divided into four groups. Proteins in group i (osteopontin, fetuin A, fetuin B, BSP, and MATN3) showed expression differences under mechanical stimulation and experienced positive selection during vertebrate evolution. Group ii proteins exhibited no positive selection and no differential expression under mechanical stimulation. None of the proteins tested belonged to group ii. The proteins in group iii (osteonectin, ALP, and TGF-β3) were under purifying selection and exhibited differences in protein expression under the static versus the mechanical stimulation conditions. The proteins in group iv (annexin A1 and RSPO1) were under purifying selection and exhibited no differences in protein expression during the mechanical stimulation experiments.

To test the conservation hypothesis of bone ECM protein mechano-responses, we have cultured osteoblast cells under static and mechanical stimulation conditions and quantified protein expression levels of ten selected bone ECM proteins. The selected proteins for the mechanical stimulation experiments included those with low, medium, and high similarity in their amino acid sequences (Fig. 1A). Seven proteins (osteonectin, alkaline phosphatase [ALP], transforming growth factor-β3 [TGF-β3], osteopontin, bone sialoprotein [BSP], annexin A1, and R-spondin 1 [RSPO1]) have similar sequence and structure conservation values. However, three poorly conserved proteins at the sequence level (MATN3, fetuin A, and fetuin B) are moderately conserved at the structure level (Fig. 1AB, table S4). Moreover, we performed an analysis of positive selection (see Materials and Methods) of all protein groups (*N* = 255) from the Phylobone database (*11*) to determine the putative effect of evolutionary transitions in mechano-skeletal adaptations to mechanical loading.

### Mechanical loading experimentsa

We speculated that highly conserved bone ECM proteins (under purifying selection) play a major role in bone homeostasis of vertebrates. In contrast, poorly conserved proteins (under positive selection in the major locomotion transitions of evolution (Fig. 1C)) are crucial to achieve mecano-skeletal adaptations in the evolution of vertebrates, including the upright bipedal locomotion in humans. Indeed, the expressions of poorly (osteopontin, BSP, fetuin A, and fetuin B) and medium (MATN3) conserved proteins (Fig. 1D) was enhanced in osteoblasts under mechanical stimulation conditions (Fig. 2 and 3). The osteoblast-like cells were confluent after 1 week and osteoblast differentiation was observed in some cells, as indicated by ALP staining (Fig. 2). The ALP-positive area increased with culture time. Under the static condition, osteoblast differentiation started after 1 week and differentiated into mature osteoblasts after 4 weeks. The cells exposed to mechanical stimulation differentiated into osteoblasts after 2 weeks and mineral deposition occurred after 4 weeks. ALP staining revealed that the mechanical stimulation enhanced osteoblast differentiation.

**Fig. 2.**
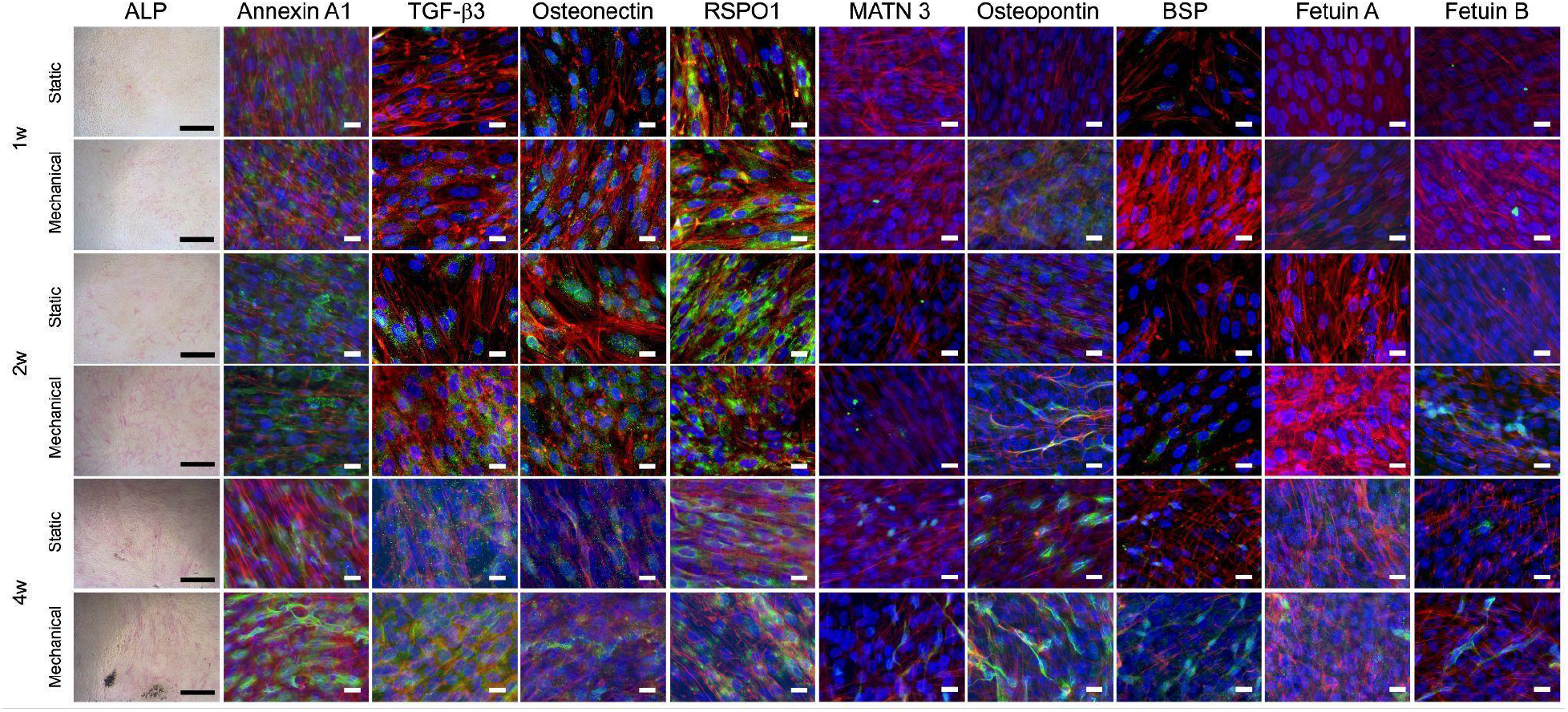
Staining of osteoblasts with or without mechanical stimulation. Osteoblasts were cultured in osteoinductive media with or without mechanical stimulation. The cells under the static condition started the osteoblast differentiation after 1 week and differentiated into mature osteoblasts after 4 weeks. The cells exposed to mechanical stimulation differentiated into osteoblasts after 2 weeks and mineral deposition occurred after 4 weeks. ALP staining revealed that the mechanical stimulation enhanced osteoblast differentiation. Scale bar for ALP images= 500μm. Immunofluorescent staining shows that the osteoblasts expressed the target proteins. Merged images of actin (red), nuclei (blue) and the targeted proteins (green) are shown. The red, green, and blue color separated images are available in fig. S1. Four proteins (RSPO1, annexin A1, osteonectin, and TGF-β3) were expressed from early to late stages of osteoblast differentiation. Two proteins (RSPO1 and annexin A1) were expressed under both the static and mechanical stimulation conditions. Five proteins (BSP, MATN3, osteopontin, fetuin A, and fetuin B) were observed as dot-like structures in the early stage of osteoblast differentiation and bundle-like structures or aggregated structures in the late stage of osteoblast differentiation. Scale bar for immunofluorescent images= 100μm

**Fig. 3.**
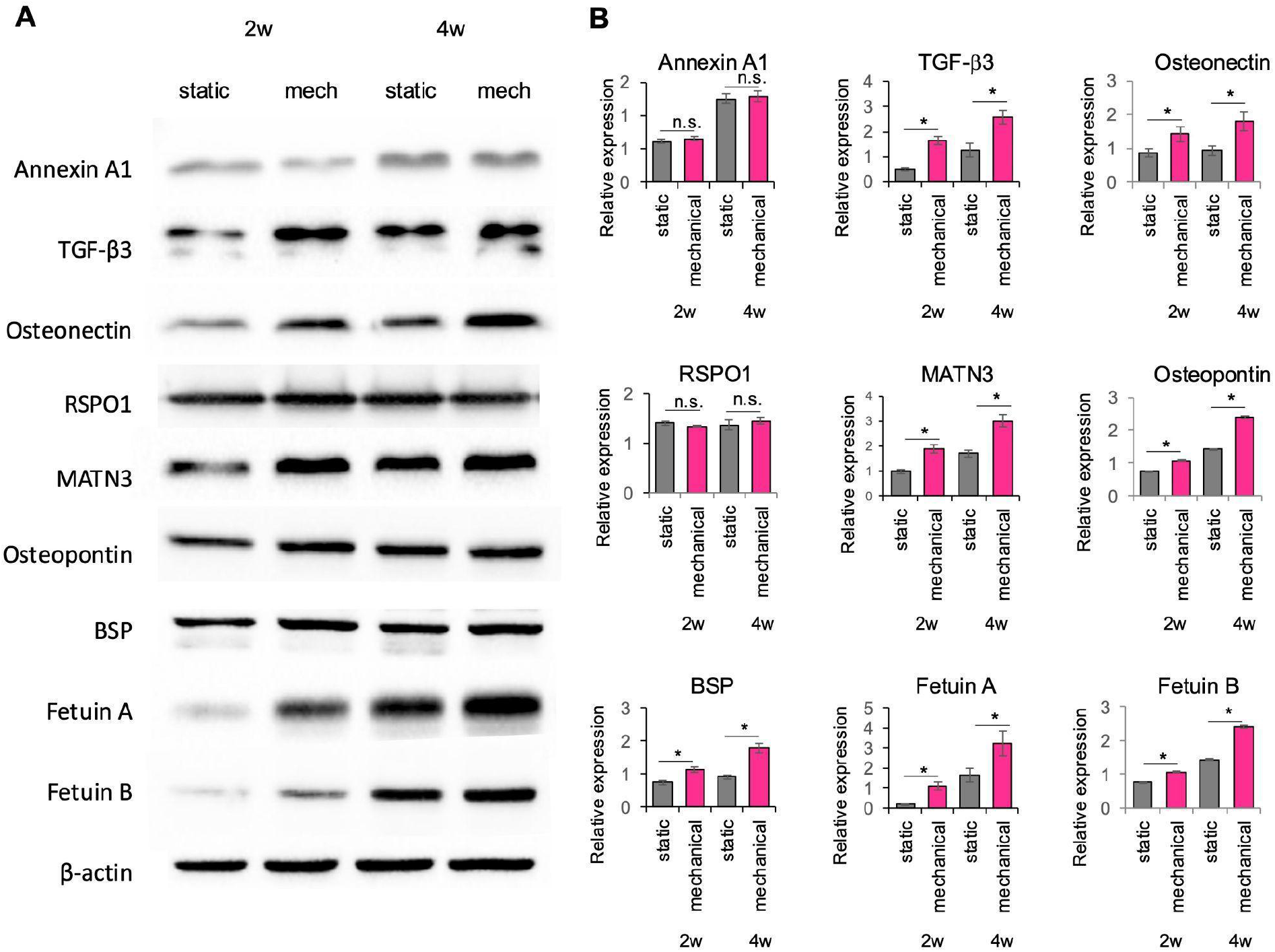
Western blot analyses of osteoblasts with or without mechanical stimulation. **(A)** Western blotting of the cell lysates of the osteoblasts cultured in osteoinductive media with or without mechanical stimulation for 2 and 4 weeks. **(B)** Densities of Western blotting bands of the target proteins normalized with β-actin. There were no significant differences in the expressions of RSPO1 and annexin A1 between the static and mechanical stimulation conditions. The expressions of the other target proteins (TGF-β3, osteonectin, MATN3, osteopontin, BSP, fetuin A, and fetuin B) was approximately 1.5 to 2-times greater under the mechanical stimulation condition compared to the static condition. As compared with the static condition, the expression of fetuin A was six times greater with mechanical stimulation for 2 weeks. Osteonectin and BSP expression increased between 2 and 4 weeks under the mechanical stimulation condition but not under the static condition. Data are reported as means ± standard deviation (SD). \**p* < 0.05 for the comparison of the static condition versus the mechanical stimulation condition.

Immunofluorescent staining showed that the osteoblasts expressed the target proteins, including annexin A1, TGF-β3, osteonectin, RSPO1, MATN3, osteopontin, BSP, fetuin A, and fetuin B (Fig. 2). Three proteins (RSPO1, annexin A1 and osteonectin) were expressed from early to late stages of osteoblast differentiation. Two proteins (RSPO1 and annexin A1) were expressed under both static and mechanical stimulation conditions. TGF-β3 expression was weak after 1 week but gradually increased after 2 weeks. Two proteins (BSP and MATN3) were seen as dot-like structures after mechanical stimulation for 2 weeks and as bundle-like structures after mechanical stimulation for 4 weeks. The expression of osteopontin was weak after 1 week under the mechanical stimulation condition and after 2 weeks under the static condition. After 4 weeks, osteopontin was seen as dot-like structures under the static condition and as bundle-like structures under the mechanical stimulation condition. The expression of fetuin A was weak after 2 weeks and increased after 4 weeks. After 4 weeks, fetuin A was seen as dot-like structures under the static condition and as aggregated structures under the mechanical stimulation condition. The expression of fetuin B was weak after 1 week of mechanical stimulation, but increased after 2 weeks under the mechanical stimulation condition. After 2 weeks and 4 weeks of mechanical stimulation, fetuin B was observed as bundle-like structures.

Western blotting revealed that there were no significant differences between the static and mechanical stimulation conditions in the expressions of RSPO1 and annexin A1 (Fig. 3). The expressions of the other target proteins, including TGF-β3, osteonectin, MATN3, osteopontin, BSP, fetuin A, and fetuin B was increased approximately 1.5 to 2-times under the mechanical stimulation condition compared to the static condition. After mechanical stimulation for 2 weeks, fetuin A expression was 6-times greater than the static condition. Osteonectin and BSP expression increased between 2 and 4 weeks under the mechanical stimulation condition but not under the static condition.

### Evolutionary adaptations

Positive selection, also known as Darwinian selection, is an evolutionary process in which advantageous mutations thrive in the population and promote the emergence of new phenotypes (*19*). The proteins under positive selection during the locomotion transition of vertebrate evolution (osteopontin, BSP, fetuin A, fetuin B, and MATN3) (Fig. 1C) exhibited increased expression during mechanical stimulation (Fig. 1D). These proteins showed a positive ratio of non-synonymous and synonymous mutations (dN/dS > 1). Notably, we could not find any protein under positive selection with no differential expression under the mechanical stimulation condition (Fig. 1D). These results are in agreement with the conservation hypothesis, i.e. proteins with a large number of cumulated adaptive mutations (positive selection) during locomotion transitions play a major role in bone mechano-responses to mechanical stimulation (Fig. 4). Our hypothesis suggests a punctuated equilibrium model for bone ECM proteins involved in bone mechano-responses that combine rapid evolutionary (locomotion transitions) and relatively steady phases.

**Fig. 4.**
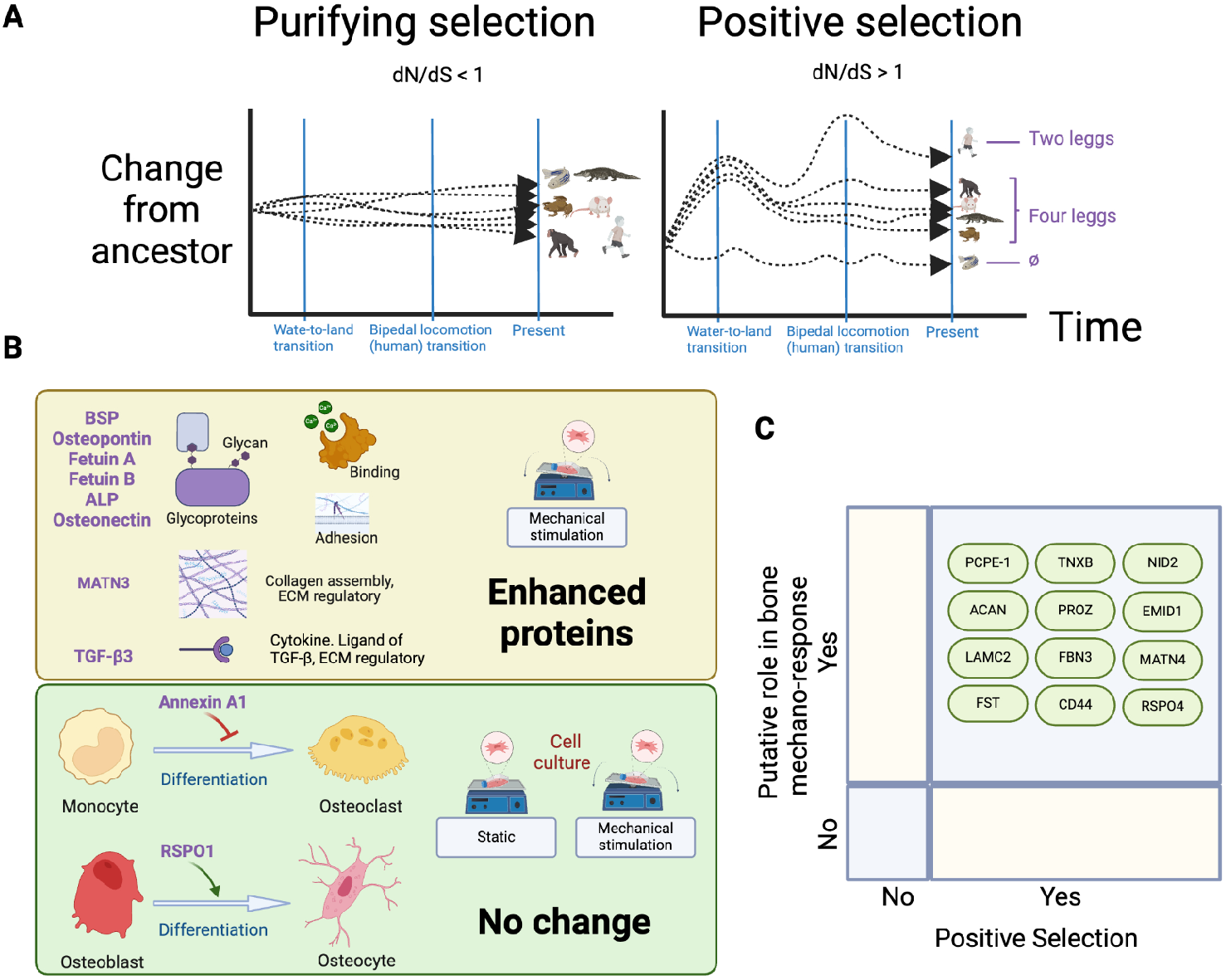
Proteins involved in bone mechano-responses. **(A)** Schematic representation of proteins under purifying (dN/dS < 1) and positive (dN/dS > 1) selection during the water-to-land transition in terrestrial vertebrates and the bipedal locomotion transition in humans. Proteins in the bone extracellular matrix (ECM) involved in mechano-response are hypothesized to accumulate adaptive mutations (shown as positive selection) during the evolutionary transitions that altered locomotion mechanisms in vertebrates. **(B)** In the mechanical stimulation experiments, the expression of eight regulatory bone ECM proteins was enhanced: six glycoproteins (BSP, osteopontin, fetuin A, fetuin B, and osteonectin) with functions involved in molecular binding (e.g., binding of calcium) and cell adhesion and two non-glycoproteins (MATN3 involved in collagen assembly, and the cytokine, TGF-β3). **(C)** Additional bone ECM proteins, that presents signs of positive selections during water-to-land and bipedalism transitions, and with functions potentially involved in bone mechano-responses (Tenascin-X, TNXB; Aggrecan core protein, ACAN; Nidogen-2, NID2; Vitamin K-dependent protein Z, PROZ; EMI domain-containing protein1, EMID1;Laminin subunit gamma-2, LAMC2; Fibrillin-3, FBN3; Matrilin-4, MATN4; Follistatin, FST; CD44 antigen, CD44; R-spondin-4, RSPO4).

The opposite of positive selection is purifying selection, also known as negative selection, a process whereby adverse mutations are eliminated from the population (*20*). Conserved bone ECM proteins throughout vertebrate evolution, under purifying selection, are expected to play a major role across vertebrates (from zebrafish to human), independently of mechanical stimulation and static conditions. The expression patterns of two proteins under purifying selection (RSPO1 and annexin A), were not significantly altered by the mechanical stimulation experiments (Fig. 1D). However, three proteins (osteonectin, ALP, TGF-β3), with dN/dS<1 between zebrafish and human (under purifying selection), showed enhanced expression during the mechanical stimulation experiments (Fig. 1D).

### Differential protein expression

The proteins under positive selection showed higher expression rates during the mechanical stimulation experiments compared to the static condition, making positive selection an evolutionary marker of their key role. However, differential protein expression rates in proteins under both positive and purifying selection arise due to genetic, genomic, epigenetic, and post-translational modification factors.

Gene duplications have contributed to the evolution of vertebrate skeleton. For example, after vertebrates evolved the mineralized skeleton, the proteins from the secretory calcium-binding phosphoprotein (such as proteins SPARC [i.e. osteonectin] and SPARCL1) originated by whole genome duplication (*21*). Gene duplications can lead to novel functions, as well as to a beneficial increase in gene expression (*22*). However, no significant differences were found between human and zebrafish, in the gene copy number of the selected proteins (table S7). Differential expression rates between the static and mechanical stimulation conditions may also have arisen due to translational selection (i.e., a differential adaptation of the protein to the translational machinery of the cell that results in higher expression rates) (*23*).Translational selection is a strong driver of evolution of vertebrate genomes that shape codon usage patterns (*24*), and it has been well described in the three domains of live (eukaryotes, bacteria and archaea) (*25, 26*). Although the selected proteins are putatively highly expressed based on codon usage biases (table S8), codon biases may not explain functional adaptations, such as cells growing under mechanical and static conditions. Furthermore, non-collagenous proteins in the bone matrix are in a highly complex environment, closely intertwined with collagen fibrils, and are differentially expressed in different anatomical locations and body tissues (table S9 and S10). Thus, further experiments beyond the scope of this article are needed to determine the role of epigenetics, post-translational modifications and protein-protein interaction factors in the gene expression of bone ECM proteins (*27*).

## Discussion

Mechanical loading increases bone mass by stimulating bone-forming osteoblasts and osteocytes through a cascade of intracellular pathways involving several biomolecules, such as nitric oxide, prostaglandins, bone morphogenetic proteins (BMP), and Wnts (*28*). Moreover, experiments in knockout mice have shown the role of an ECM protein (osteopontin) in response to the mechanical stimulation (*9*). In this study, we identified eight proteins that exhibited higher expression during mechanical stimulation (ALP, BSP, fetuin A, fetuin B, MATN3, osteonectin, osteopontin, and TGF-β3) and two proteins with no differential expression in mechanically stimulated cell cultures (annexin A1 and RSPO1). Proteins in the bone ECM can be classified into structural and regulatory roles (*11*). Additionally, regulatory proteins can be further divided into those that regulate bone matrix structure and those that control bone cell growth and differentiation. In our experiments, seven proteins that exhibited a higher expression during mechanical stimulation are involved in the regulation of the bone matrix, with functions that encompass glycoproteins (BSP, osteopontin, osteonectin, ALP, fetuin A, and fetuin B), calcium-binding proteins (BSP), and collagen assembly regulation (MATN3) (Fig. 4B, table S11-12). In addition, the expression of the cytokine TGF-β3, which regulates the early stages of osteoblast differentiation (*29*), was enhanced by mechanical stimulation. However, the proteins involved in the regulation of osteoclast differentiation (annexin A1) and osteoblast differentiation to osteocytes (RSPO1) were not enhanced by mechanical stimulation.

The water-to-land transition in vertebrates involved the evolution of the skeletal system to support locomotive activity in the terrestrial environment (*30*). Such the evolution of the skeletal system has been linked with immune molecules, for example, the receptor activator of the nuclear factor-κB ligand (RANKL)–RANK system has been conserved since the emergence of the adaptive immune system with T and B cells in cartilaginous fish (*30*). Furthermore, during the evolution to bipedalism in humans, the skeletal system was adapted for walking regularly in an upright position (*16, 17*). This involved molecular changes in the BMP (*31*). In agreement with the conservation hypothesis presented in this study, functionally relevant proteins can be predicted by observing changes during the two significant locomotion transitions in the evolution of vertebrates (Fig 4A). Fetuin A showed the most striking change in protein expression during mechanical stimulation, with the immunofluorescent staining and Western blotting revealing that its expression was enhanced in osteoblasts by mechanical stimulation (Fig. 2 and 3). In agreement with the conservation hypothesis, fetuin A exhibits strong signs of positive selection during the water-to-land transition of vertebrates and the bipedalism transition in humans. Furthermore, fetuin A protein shows low amino acid conservation (Fig. 1A, table S2) and medium structural conservation (Fig. 1B, table S4). Fetuin A is known to influence the production of inflammation mediators, inhibit ectopic calcification in soft tissue (*32*), and reduce biomaterial particle-induced osteolysis (*33*). Therefore, Fetuin A is a strong candidate as a biomarker for skeletal disorders like osteoporosis and rheumatoid arthritis, and a potential target for drug treatment.

The first-line of osteoporosis treatments are bisphosphonates and the monoclonal RANKL antibodies. The targets of these drugs are calcium minerals in bone ECM and RANK receptors on osteoclast precursors (*34*), respectively. Among the selected proteins, fetuin A is potentially the strongest candidate for further exploration. However, additional proteins involved in mechanical loading (ALP, BSP, fetuin B, MATN3, osteonectin, osteopontin, and TGF-β3) could be potential drug targets for the treatment and prevention of osteoporosis. Moreover, other proteins of the bone ECM that play a regulatory role, such as those with roles in osteogenesis and bone degradation (*35*), may be targets for new treatments for bone regeneration (*11*). Furthermore, we speculate that ECM proteins with signs of positive selection during the water-to-land and bipedalism transitions, with regulatory functions in the bone matrix (Fig. 4C), may play a significant role in the bone mechano-response to mechanical loading.

## Materials and Methods

### Protein sequences

Protein sequences were collected from the Phylobone database (*11*). This database contains a list of 255 putative bone exctracellular matrix (ECM) proteins found in 31 species of vertebrates, including the species selected for this study. We selected one representative species from each of the major taxonomic groups of vertebrates: bony fishes (*D. rerio*), amphibians (*Xenopus laevis*), reptiles (*Alligator sinensis*), rodents (*Mus musculus*), and primates (*Pan troglodytes* and *H. sapiens*).

### Protein sequence similarity (percentage of identity between D. rerio and H. sapiens)

All protein homologs from *H. sapiens* and *D. rerio* were compared. Sequence pairs were aligned using the program Muscle (*36*) and an in-house script was utilized to calculate the percentage of identity between pairs of proteins (i.e. the number of amino acid matches and mismatches between *H. sapiens and D. rerio*). In cases where the protein sequences of *D. rerio* were not found (e.g., BSP), we assigned 0.0% of identity percentage.

### Selection of proteins for mechanical stimulation experiments

Ten bone ECM proteins were selected for the mechanical stimulation experiments (table S13 and Fig. 1A) based on the following: 1) the length of the sequences (proteins between 250 and 1,000 amino acids long); 2) percentage of identity (low: < 33.0%, medium: ≥ 33.0–66.0%, and high: ≥ 66.0%); and 3) putative functional importance of these proteins in bone ECM based on the literature (*11*). The ten proteins selected were: 1) five proteins, with less than 30% of protein identity were selected (Osteopontin, Fetuin A, Fetuin B, Bone sialoprotein (BSP), and R-spondin 1 (RSPO1)); 2) one moderately conserved protein was selected (Matrilin 3 (MATN3)); and 3) three highly conserved proteins (osteonectin (also known as secreted protein acidic and rich in cysteine, SPARC), annexin A1, alkaline phosphatase (ALP), and transforming growth factor β3 (TGF-β3)).

### Pairwise structure alignment

All proteins from *H. sapiens* were compared with their homologs in *D. rerio* (when the sequence was available) using a pairwise structure alignment tool to align the 3D structure of the protein (*37*). Predictions of protein structures were based on artificial intelligence AlphaFold models (*38*). The Root Mean Square deviation (RMSD) and the TM-Score were used to quantify similarities between the protein structures (*39*).

### Preparation of sequences for the analysis of positive selection

Multiple sequence alignments (MSAs) were performed and phylogenetic trees were reconstructed using the server NGPhylogeny (*40*). This server aligns proteins (raw MSAs) using the program MAFFT (*41*), and removes non-conserved regions in the alignment (clean MSAs) with the program Block Mapping and Gathering with Entropy (BMGE) (*42*). The Clean MSAs are the used for the reconstruction of phylogenetic trees using the program PhyML (*43*). The raw MSAs together with the corresponding coding sequences of DNA, were used as input for the server PAL2NAL (*44*) to convert the raw MSAs of proteins to the corresponding DNA sequence alignment of codon alignment (codon MSAs).The resulting codon MSAs were used for the calculation of non-synonymous (dN) and synonymous (dS) substitution rates.

### Codon pairwise alignments

Pairwise alignments of protein sequences were performed with the Sequence Manipulation server (*45*). The Smith-Waterman algorithm and the BLOSUM62 matrix were used to perform the pairwise alignments. These protein alignments were converted into codon pairwise alignments using the program PAL2NAL (*44*) and used for the calculation of dN and dS ratios.

### Prediction of positive selection with dN and dS ratios

Predictions of positive selection were estimated with the dN/dS ratio, which is the ratio of the number of non-synonymous substitutions per non-synonymous site (pN) to synonymous substitutions per synonymous site (pS) (*46*). We have analyzed the dN/dS ratio in five vertebrate species (table S1) using an indel-aware algorithm (https://github.com/clarabrunet/iDNDS) that analyzes the ratio at different levels of the species tree and uses an indel-aware table of expected synonymous and non-synonymous mutations (table S14). We considered proteins to be under positive selection when dN/dS was greater than 1.

### Cell culture

Osteoblast-like cells (MG-63 cell line) were seeded into 12-well cell culture plates at a density of 2×10^4^ cells/well and cultured in Dulbecco’s modified Eagle’s medium (DMEM; Gibco 21969-035, NY, USA) supplemented with 10% fetal bovine serum (Gibco 10270-098, NY, USA), _L_-glutamine (Gibco 25030081, NY, USA), and 100 U/ml penicillin-streptomycin (Gibco 15140-122, NY, USA) in a humidified atmosphere of 5% CO_2_ in air at 37°C. After reaching confluence, the cells were cultured in osteoinductive medium including 100 nM dexamethasone (Sigma-Aldrich, St. Louis, MO, USA), 10 mM β-glycerophosphate (Sigma-Aldrich, St. Louis, MO, USA) and 50 µg/ml ascorbic acid (Sigma-Aldrich, St. Louis, MO, USA). The cell culture plates were placed on a see-saw rocker for 30 min/day to induce mechanical stimulation (*47*).

After the culture for 1, 2 and 4 weeks, the cells were fixed in 4% paraformaldehyde for 20 min. To visualize the differentiation into osteoblasts, the cells were stained with alkaline phosphatase (ALP) (Sigma-Aldrich, St. Louis, MO, USA) and observed using a microscope (Olympus IX71).

### Immunofluorescent staining

After the cell culture for 1, 2 and 4 weeks, the cells were fixed with 4% paraformaldehyde for 20 min. The cells were incubated in a solution for blocking and permealization solution including 0.2% Tween 20 and 1% bovine serum albumin (BSA) in phosphate-buffered saline (PBS). Anti-human primary antibodies (osteonectin, PA578178; RSPO1, PA5121183; TGF β-3, PA599186; annexin A1, 713400; MATN3, PA520727; fetuin A. PA551594; fetuin B, PA529468; BSP, Bs-2668R; osteopontin, 229521AP, Invitrogen, CA, USA) diluted in PBS including 0.2% BSA were added for 1h at room temperature. Following PBS washes, the cells were incubated in a secondary antibody (Alexa Fluor ™ 488 Goat anti-rabbit IgG antibody, A11008, CA, USA) diluted in PBS including 0.2% BSA. The cells were stained for rhodamine phalloidin (Invitrogen R415, CA, USA) and Hoechst 33258 (Sigma-Aldrich, 94403, St. Louis, MO, USA). The fluorescent signals were observed using a fluorescence microscope (Olympus IX71).

### Western blotting

After the cell culture for 2 and 4 weeks, the cells were lysed with 300µl of lysis buffer containing 50mM Tris-HCl (pH6), 150mM NaCl and 1% Triton X-100. The cells were scraped, collected into tubes and sonicated for 10 seconds. The cell lysates were frozen at -80 °C until use.

The protein samples were mixed with 10x Sodium dodecyl-sulfate (SDS) loading buffer (Bio-rad 1610747, CA, USA) and heated at 95 °C for 5 min. The proteins samples were electrophoresed at 60V for 15 min and further 120V for 120 min on 5% stacking gel with 10% (for BSP, Fetuin A, Fetuin B, and MATN 3) or 12% (for osteonectin, RSPO1, TGF-β3, annexin A1, osteopontin, and β-actin) SDS polyacrylamide. The proteins were transformed from SDS gel to nitrocellulose membrane at 35V for 30 min. The nitrocellulose membrane was incubated in blocking solution including 5% skim milk in TBST (Tris-buffered saline including 0.1% Tween 20) for 60 min at room temperature with shaking. After washing the nitrocellulose membrane in TBST for 10 min at room temperature with shaking three times, the membrane was incubated in blocking solution including primary antibodies at a 1:500 dilution at 4 °C overnight. Goat anti-rabbit IgG horseradish peroxidase (HRP)-conjugated secondary antibody (Invitrogen 31460, CA, USA) were diluted in blocking solution and added to the membrane at room temperature for 60 min. After washing with TBST, the protein bands were visualized using a chemiluminescence reagent (Thermo Scientific™ 34585, MA, USA), captured with a ChemiDoc (Bio-Rad, CA, USA) and quantified using Image J software (version 1.41, National Institute of Health).

### Statistical analysis

Accurate quantifications of the different samples in cell study were achieved by performing four independent experiments. Statistical analysis across the experimental groups was performed using the analysis of variance (ANOVA) with Tukey’s post-hoc test for multiple comparisons, using the SPSS software package (version 29, Chicago, IL, USA). The statistically significant level was set at *p* < 0.05 for all the tests. All data has been expressed as mean ± standard deviation (SD).

## Acknowledgments

This study was supported by the IT-CSC Finland (Project ID #2004931). We thank Erasmus+ undergraduate students Ms. Valentine Jarry and Ms. Aude I Sanchez for their help in performing the mechanical loading experiments and other members and collaborators of the Phylobone project for their helpful discussions. We thank Ms. Samira El Manfi from Prof. Ulvi Gursoy’s group and Ms. Katja Sampalahti, Ms. Tatjana Peskova, and Dr. Vuokko Loimaranta for their technical assistance in performing the Western blotting.

## Funding

Sigrid Juselius Foundation 230131 (MN, PP)

JSPS Grants-in-Aid for Scientific Research JP23K08670 (MN, PP) Murata Foundation M20-072 (MN)

Turku Collegium for Science, Medicine and Technology (MN)

## Author contributions

Conceptualization: MN, PP

Methodology: MN, PP, SS

Investigation: MN, PP, SS, KH, MFR,

CB Visualization: MN, PP

Funding acquisition: MN, PP

Project administration: MN, PP Supervision: MN, PP

Writing – original draft: MN, PP, SS, CB Writing – review & editing: MN, PP

## Data and materials availability

The authors declare that all data supporting the findings of this study are available within this article and its supplementary information files. Bone ECM proteins are available in the Phylobone database at https://phylobone.com. The algorithm implemented to calculate dN/dS ratios is available at https://github.com/clarabrunet/iDNDS.

## Supplementary Materials

Fig. S1

Tables S1 to S14

References (*1*–*66*)

