## Supplementary material for "Bone mechano-response is driven by locomotion transitions during vertebrate evolution"

**The PDF file includes:**

Fig. S1
Tables S1 to S14
References

### Supplementary Figures

**A** Annexin A1

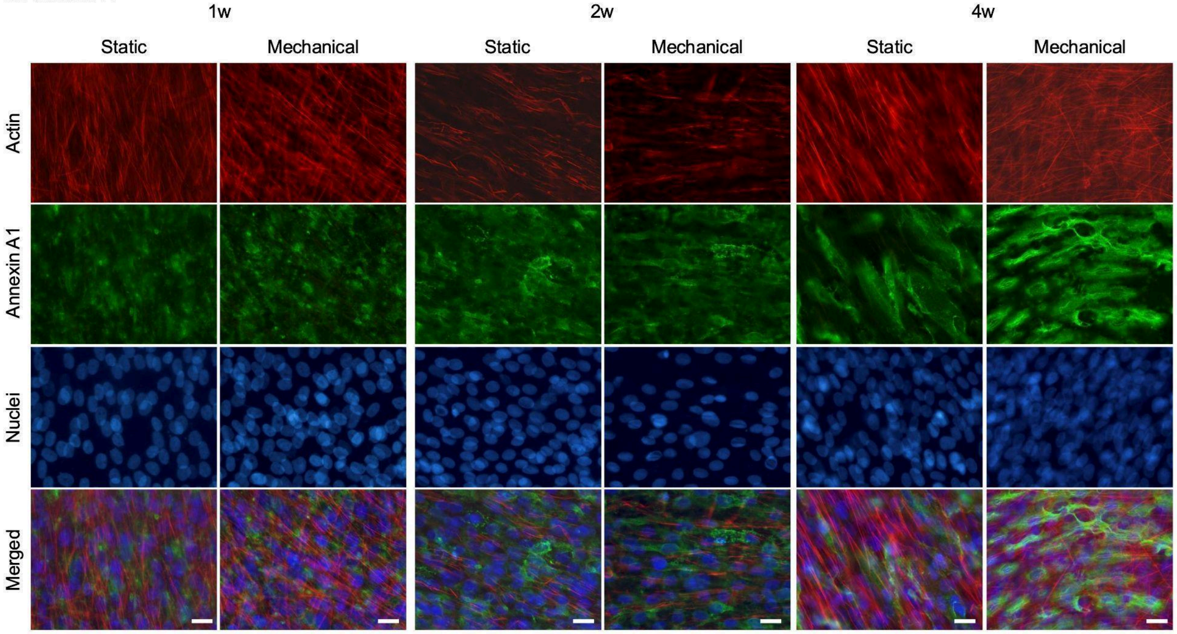

**B** TGF- β3

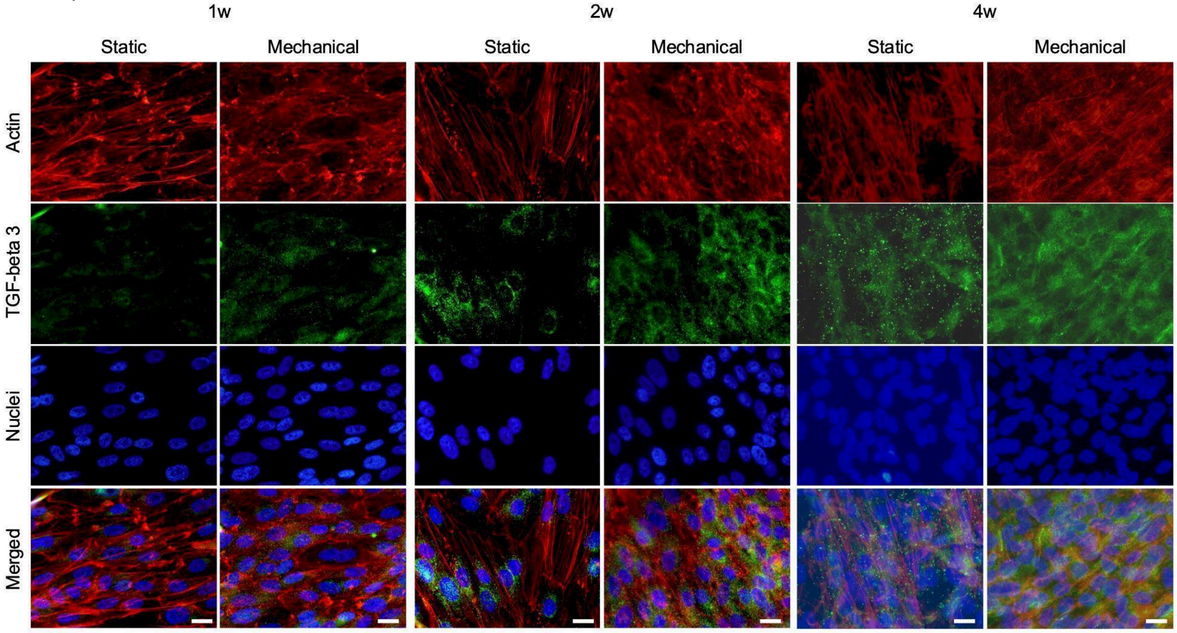

**C** Osteonectin

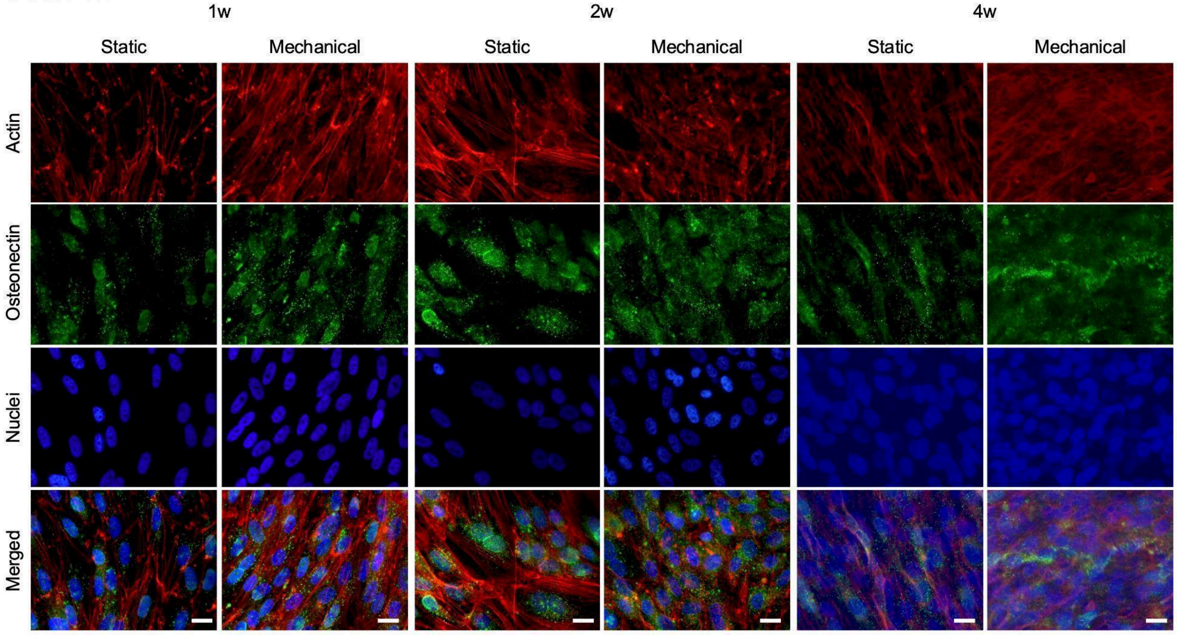

**D** RSPO1

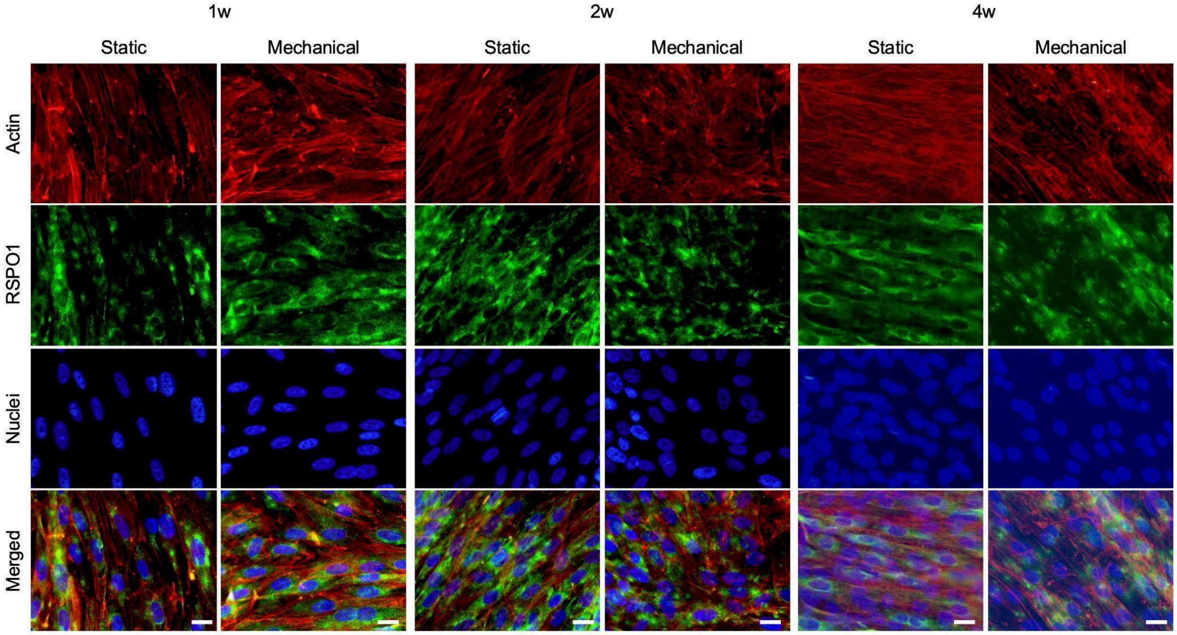

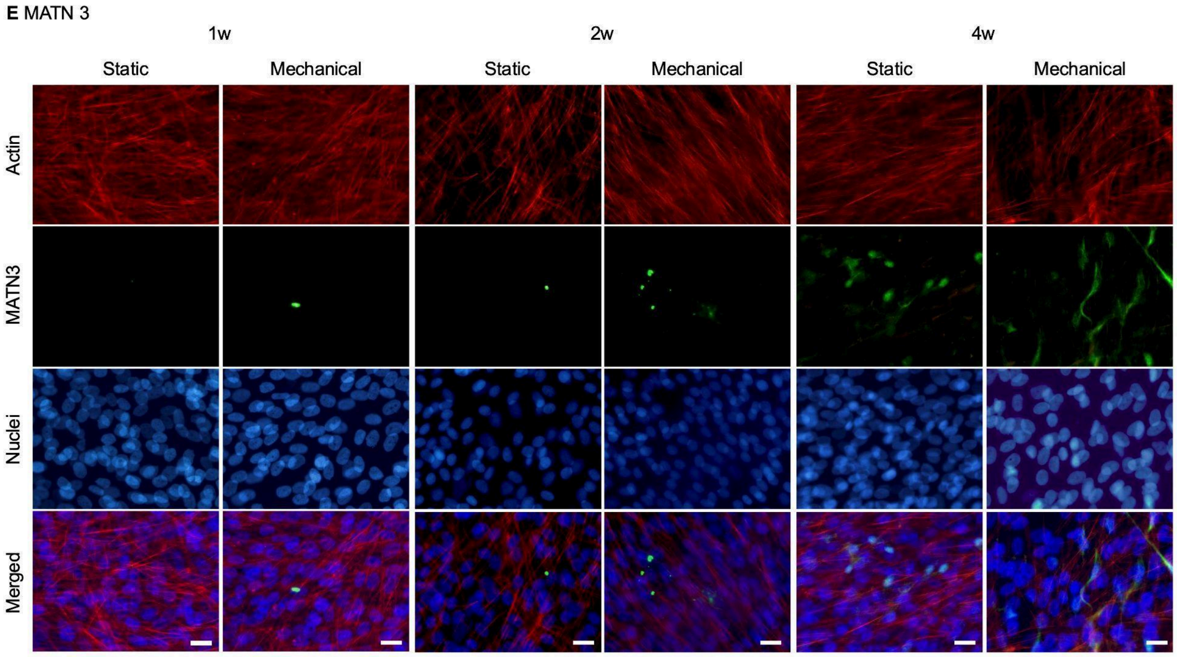
E MATN 3
1w
2w
4w
Static
Mechanical
Static
Mechanical
Static
Mechanical
Actin
MATN3
Nuclei
Merged

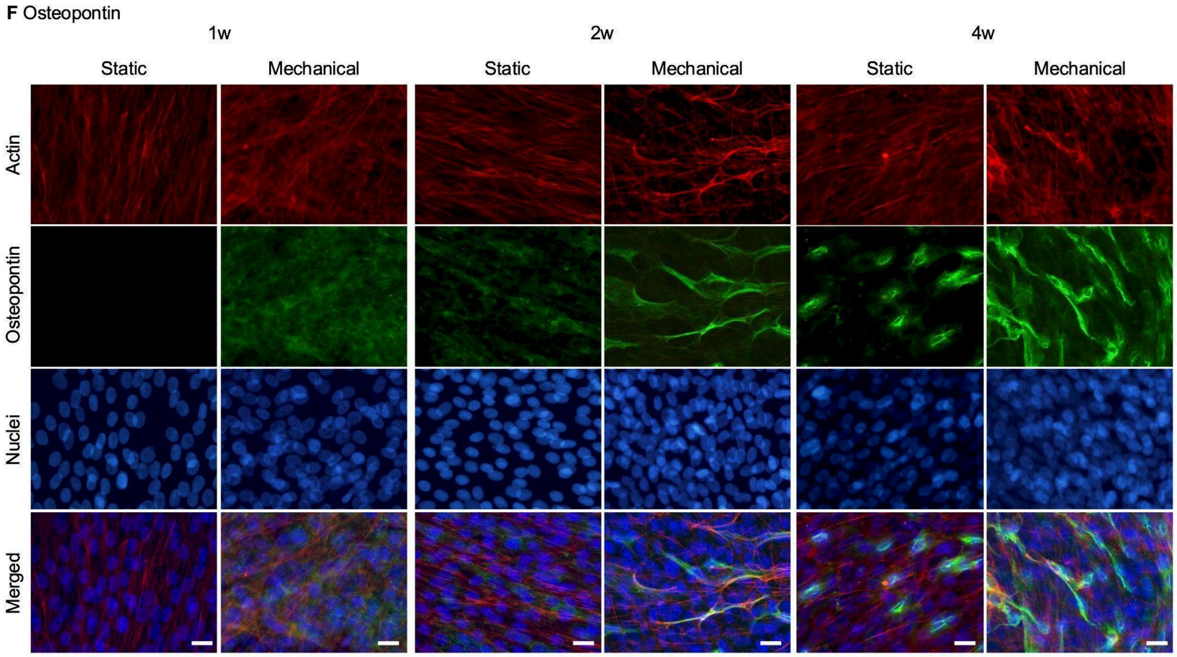
F Osteopontin
1w
2w
4w
Static
Mechanical
Static
Mechanical
Static
Mechanical
Actin
Osteopontin
Nuclei
Merged

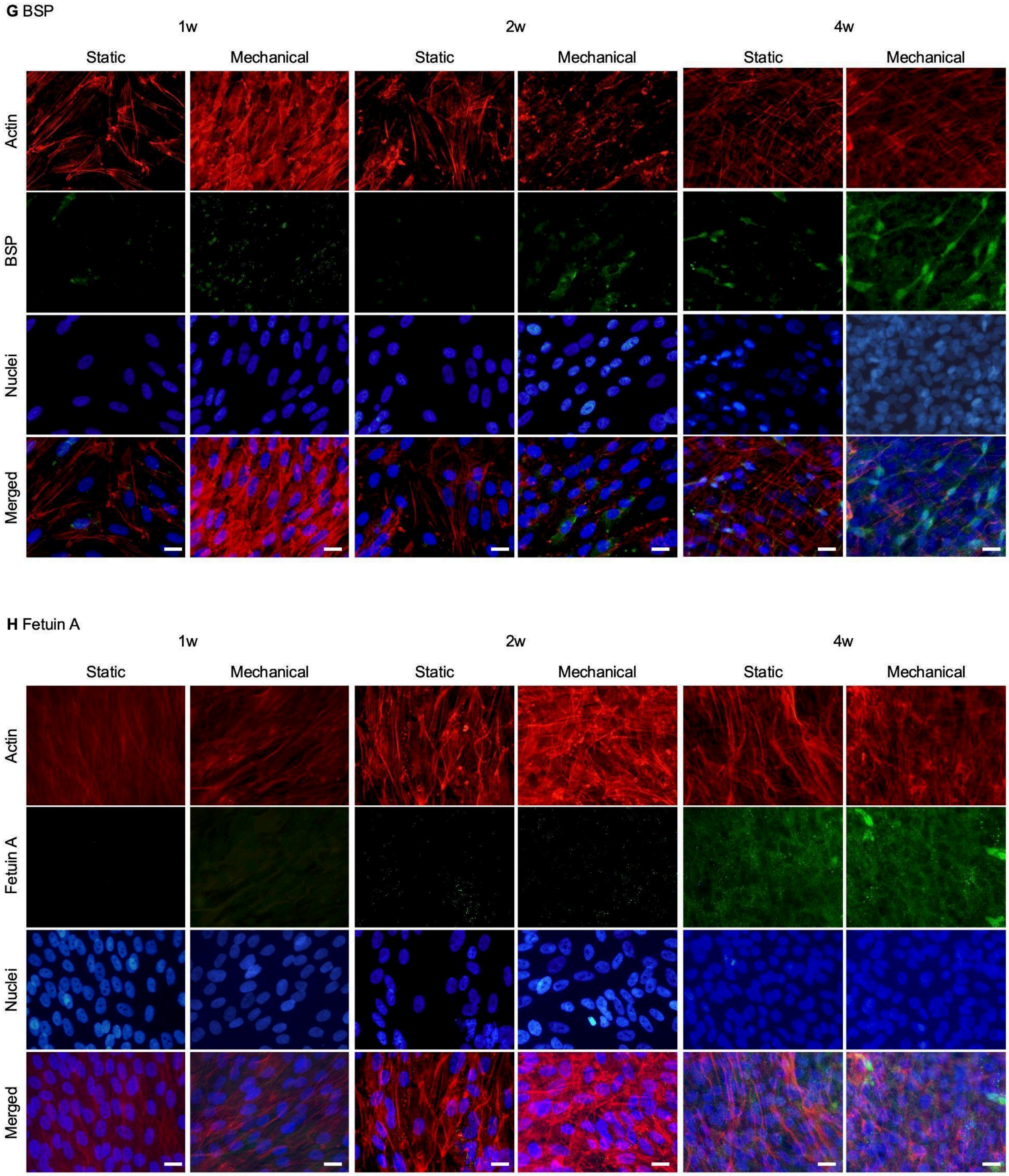

**Fig. S1.**
Immunofluorescent staining of osteoblast-like cells cultured in osteoinductive media with or without mechanical stimulation for 1, 2, and 4 weeks. The merged images of actin (red), nuclei (blue) and the targeted proteins (green) are shown. The osteoblasts expressed the target proteins such as (A) annexin A1, (B) TGF-β3, (C) osteonectin, (D) RSPO1, (E) MATN3, (F) osteopontin, (G) BSP, (H) fetuin A, and (I) fetuin B. Scale bar =100mm.

#### Supplementary Tables

**Table S1.**
List of vertebrate species used in this study.

| Species | Common Name | Taxonomy | Proteins |
|---|---|---|---|
| *Danio rerio* | Zebrafish | Bony fishes (TAXID:7955) | 282 (in 251 groups) |
| *Xenopus laevis* | Frog | Amphibians (TAXID:8355) | 245 (in 245 groups) |
| *Alligator sinensis* | Reptile | Reptiles (TAXID:38654) | 238 (in 238 groups) |
| *Mus musculus* | Mouse | Rodents (TAXID:10090) | 253 (in 253 groups) |
| *Pan troglodytes* | Chimpanzee | Primates (TAXID:9598) | 256 (in 255 groups) |
| *Homo sapiens* | Human | Primates (TAXID:9606) | 449 (in 255 groups) |

*Proteins: Number of proteins in the bone extracellular matrix (1)*

**Table S2.**
Sequence similarity of bone extracellular matrix (ECM) proteins between *Danio rerio* and *Homo sapiens*

| Phylobone Code | Length | Match | Mismatch | Identity percentage |
|---|---|---|---|---|
| PB0210 (*) | 273 | 0 | 273 | 0% |
| PB0230 (*) | 1466 | 0 | 1466 | 0% |
| PB0249 (*) | 317 | 0 | 317 | 0% |
| PB0250 | 205 | 0 | 205 | 0% |
| PB0063 | 4262 | 510 | 3752 | 12% |
| PB0047 | 141 | 21 | 120 | 15% |
| PB0034 | 611 | 94 | 517 | 15% |
| PB0117 | 1193 | 214 | 979 | 18% |
| PB0102 | 344 | 65 | 279 | 19% |
| PB0109 | 221 | 44 | 177 | 20% |
| PB0169 | 5435 | 1102 | 4333 | 20% |
| PB0049 | 404 | 86 | 318 | 21% |
| PB0089 | 218 | 49 | 169 | 23% |
| PB0086 | 268 | 62 | 206 | 23% |
| PB0008 | 257 | 61 | 196 | 24% |
| PB0101 | 2460 | 597 | 1863 | 24% |
| PB0041 | 712 | 179 | 533 | 25% |
| PB0085 | 507 | 133 | 374 | 26% |
| PB0199 | 446 | 120 | 326 | 27% |
| PB0088 | 103 | 28 | 75 | 27% |
| PB0218 | 151 | 41 | 110 | 27% |
| PB0004 | 113 | 31 | 82 | 27% |
| PB0001 | 418 | 115 | 303 | 28% |
| PB0195 | 489 | 137 | 352 | 28% |
| PB0217 | 188 | 53 | 135 | 28% |
| PB0042 | 286 | 84 | 202 | 29% |
| PB0177 | 449 | 136 | 313 | 30% |
| PB0083 | 6344 | 1934 | 4410 | 31% |
| PB0181 | 276 | 86 | 190 | 31% |
| PB0162 | 606 | 191 | 415 | 32% |
| PB0118 | 854 | 273 | 581 | 32% |
| PB0012 | 114 | 37 | 77 | 33% |
| PB0087 | 88 | 29 | 59 | 33% |
| PB0104 | 461 | 153 | 308 | 33% |

| | | | | |
|---|---|---|---|---|
| PB0014 | 272 | 91 | 181 | 34% |
| PB0179 | 317 | 107 | 210 | 34% |
| PB0068 | 2558 | 866 | 1692 | 34% |
| PB0093 | 1353 | 469 | 884 | 35% |
| PB0154 | 1916 | 666 | 1250 | 35% |
| PB0232 | 870 | 304 | 566 | 35% |
| PB0165 | 517 | 181 | 336 | 35% |
| PB0043 | 1302 | 460 | 842 | 35% |
| PB0013 | 266 | 96 | 170 | 36% |
| PB0189 | 155 | 56 | 99 | 36% |
| PB0153 | 1675 | 627 | 1048 | 37% |
| PB0071 | 1685 | 636 | 1049 | 38% |
| PB0080 | 1733 | 654 | 1079 | 38% |
| PB0056 | 248 | 95 | 153 | 38% |
| PB0159 | 784 | 302 | 482 | 39% |
| PB0222 | 1498 | 576 | 922 | 39% |
| PB0175 | 437 | 169 | 268 | 39% |
| PB0240 | 384 | 149 | 235 | 39% |
| PB0039 | 337 | 132 | 205 | 39% |
| PB0009 | 422 | 166 | 256 | 39% |
| PB0045 | 800 | 318 | 482 | 40% |
| PB0076 | 988 | 393 | 595 | 40% |
| PB0090 | 2892 | 1152 | 1740 | 40% |
| PB0180 | 347 | 139 | 208 | 40% |
| PB0255 | 252 | 102 | 150 | 41% |
| PB0058 | 722 | 294 | 428 | 41% |
| PB0016 | 250 | 103 | 147 | 41% |
| PB0194 | 951 | 392 | 559 | 41% |
| PB0060 | 261 | 108 | 153 | 41% |
| PB0116 | 532 | 222 | 310 | 42% |
| PB0002 | 119 | 50 | 69 | 42% |
| PB0136 | 571 | 240 | 331 | 42% |
| PB0193 | 532 | 224 | 308 | 42% |
| PB0070 | 1654 | 707 | 947 | 43% |
| PB0077 | 4654 | 1989 | 2665 | 43% |
| PB0141 | 1919 | 822 | 1097 | 43% |
| PB0132 | 405 | 175 | 230 | 43% |
| PB0110 | 473 | 205 | 268 | 43% |

| | | | | |
|---|---|---|---|---|
| PB0105 | 1859 | 808 | 1051 | 44% |
| PB0155 | 344 | 151 | 193 | 44% |
| PB0097 | 1802 | 795 | 1007 | 44% |
| PB0152 | 4417 | 1947 | 2470 | 44% |
| PB0026 | 475 | 210 | 265 | 44% |
| PB0061 | 329 | 147 | 182 | 45% |
| PB0035 | 464 | 208 | 256 | 45% |
| PB0149 | 382 | 171 | 211 | 45% |
| PB0254 | 341 | 153 | 188 | 45% |
| PB0219 | 327 | 148 | 179 | 45% |
| PB0212 | 206 | 95 | 111 | 46% |
| PB0166 | 267 | 124 | 143 | 46% |
| PB0124 | 512 | 238 | 274 | 47% |
| PB0200 | 514 | 239 | 275 | 47% |
| PB0242 | 150 | 70 | 80 | 47% |
| PB0111 | 1371 | 644 | 727 | 47% |
| PB0148 | 764 | 360 | 404 | 47% |
| PB0066 | 740 | 349 | 391 | 47% |
| PB0134 | 797 | 383 | 414 | 48% |
| PB0114 | 679 | 331 | 348 | 49% |
| PB0143 | 452 | 222 | 230 | 49% |
| PB0075 | 1255 | 620 | 635 | 49% |
| PB0073 | 91 | 45 | 46 | 50% |
| PB0123 | 3720 | 1840 | 1880 | 50% |
| PB0172 | 188 | 94 | 94 | 50% |
| PB0206 | 138 | 69 | 69 | 50% |
| PB0252 | 272 | 136 | 136 | 50% |
| PB0095 | 442 | 222 | 220 | 50% |
| PB0082 | 620 | 312 | 308 | 50% |
| PB0098 | 376 | 189 | 187 | 50% |
| PB0120 | 685 | 345 | 340 | 50% |
| PB0094 | 1259 | 636 | 623 | 51% |
| PB0037 | 932 | 472 | 460 | 51% |
| PB0113 | 1669 | 851 | 818 | 51% |
| PB0051 | 3648 | 1864 | 1784 | 51% |
| PB0067 | 515 | 264 | 251 | 51% |
| PB0187 | 670 | 344 | 326 | 51% |
| PB0038 | 276 | 142 | 134 | 51% |

| | | | | |
|---|---|---|---|---|
| PB0052 | 957 | 492 | 465 | 51% |
| PB0072 | 1225 | 631 | 594 | 52% |
| PB0003 | 651 | 336 | 315 | 52% |
| PB0112 | 479 | 247 | 232 | 52% |
| PB0029 | 491 | 254 | 237 | 52% |
| PB0091 | 1681 | 870 | 811 | 52% |
| PB0244 | 613 | 318 | 295 | 52% |
| PB0140 | 1743 | 912 | 831 | 52% |
| PB0208 | 502 | 263 | 239 | 52% |
| PB0227 | 268 | 141 | 127 | 53% |
| PB0074 | 465 | 245 | 220 | 53% |
| PB0126 | 289 | 153 | 136 | 53% |
| PB0147 | 1730 | 918 | 812 | 53% |
| PB0190 | 408 | 218 | 190 | 53% |
| PB0062 | 340 | 182 | 158 | 54% |
| PB0023 | 474 | 254 | 220 | 54% |
| PB0017 | 143 | 77 | 66 | 54% |
| PB0215 | 410 | 221 | 189 | 54% |
| PB0054 | 930 | 502 | 428 | 54% |
| PB0137 | 682 | 369 | 313 | 54% |
| PB0207 | 357 | 193 | 164 | 54% |
| PB0106 | 168 | 91 | 77 | 54% |
| PB0096 | 1034 | 563 | 471 | 54% |
| PB0064 | 202 | 110 | 92 | 55% |
| PB0224 | 365 | 199 | 166 | 55% |
| PB0234 | 1074 | 586 | 488 | 55% |
| PB0010 | 177 | 97 | 80 | 55% |
| PB0025 | 2538 | 1394 | 1144 | 55% |
| PB0144 | 237 | 130 | 107 | 55% |
| PB0185 | 386 | 213 | 173 | 55% |
| PB0197 | 356 | 198 | 158 | 56% |
| PB0050 | 559 | 312 | 247 | 56% |
| PB0115 | 378 | 211 | 167 | 56% |
| PB0241 | 933 | 521 | 412 | 56% |
| PB0167 | 662 | 370 | 292 | 56% |
| PB0238 | 829 | 464 | 365 | 56% |
| PB0142 | 2077 | 1171 | 906 | 56% |
| PB0138 | 385 | 218 | 167 | 57% |

| | | | | |
|---|---|---|---|---|
| PB0191 | 201 | 114 | 87 | 57% |
| PB0216 | 271 | 154 | 117 | 57% |
| PB0107 | 716 | 408 | 308 | 57% |
| PB0214 | 456 | 260 | 196 | 57% |
| PB0048 | 556 | 322 | 234 | 58% |
| PB0103 | 315 | 183 | 132 | 58% |
| PB0121 | 2227 | 1306 | 921 | 59% |
| PB0127 | 302 | 178 | 124 | 59% |
| PB0021 | 217 | 129 | 88 | 59% |
| PB0040 | 752 | 447 | 305 | 59% |
| PB0046 | 677 | 402 | 275 | 59% |
| PB0007 | 363 | 216 | 147 | 60% |
| PB0163 | 1879 | 1120 | 759 | 60% |
| PB0235 | 1055 | 629 | 426 | 60% |
| PB0229 | 1828 | 1096 | 732 | 60% |
| PB0231 | 707 | 425 | 282 | 60% |
| PB0032 | 314 | 189 | 125 | 60% |
| PB0030 | 1751 | 1060 | 691 | 61% |
| PB0081 | 1026 | 622 | 404 | 61% |
| PB0100 | 335 | 203 | 132 | 61% |
| PB0203 | 269 | 163 | 106 | 61% |
| PB0069 | 902 | 548 | 354 | 61% |
| PB0125 | 468 | 285 | 183 | 61% |
| PB0176 | 690 | 424 | 266 | 61% |
| PB0119 | 322 | 198 | 124 | 62% |
| PB0220 | 343 | 211 | 132 | 62% |
| PB0084 | 1511 | 938 | 573 | 62% |
| PB0099 | 1187 | 737 | 450 | 62% |
| PB0015 | 429 | 270 | 159 | 63% |
| PB0221 | 350 | 220 | 130 | 63% |
| PB0020 | 502 | 317 | 185 | 63% |
| PB0158 | 496 | 313 | 183 | 63% |
| PB0135 | 3351 | 2120 | 1231 | 63% |
| PB0139 | 2267 | 1451 | 816 | 64% |
| PB0213 | 184 | 118 | 66 | 64% |
| PB0211 | 321 | 206 | 115 | 64% |
| PB0079 | 1904 | 1234 | 670 | 65% |
| PB0033 | 339 | 220 | 119 | 65% |

| | | | | |
|---|---|---|---|---|
| PB0053 | 1821 | 1183 | 638 | 65% |
| PB0028 | 318 | 207 | 111 | 65% |
| PB0174 | 248 | 162 | 86 | 65% |
| PB0182 | 402 | 263 | 139 | 65% |
| PB0145 | 684 | 449 | 235 | 66% |
| PB0198 | 375 | 246 | 129 | 66% |
| PB0044 | 1695 | 1113 | 582 | 66% |
| PB0164 | 685 | 451 | 234 | 66% |
| PB0228 | 788 | 520 | 268 | 66% |
| PB0183 | 513 | 339 | 174 | 66% |
| PB0156 | 2926 | 1938 | 988 | 66% |
| PB0171 | 454 | 301 | 153 | 66% |
| PB0226 | 700 | 465 | 235 | 66% |
| PB0146 | 978 | 662 | 316 | 68% |
| PB0178 | 378 | 256 | 122 | 68% |
| PB0170 | 2881 | 1958 | 923 | 68% |
| PB0186 | 692 | 471 | 221 | 68% |
| PB0133 | 183 | 125 | 58 | 68% |
| PB0024 | 450 | 308 | 142 | 68% |
| PB0092 | 564 | 386 | 178 | 68% |
| PB0184 | 1609 | 1101 | 508 | 68% |
| PB0031 | 497 | 341 | 156 | 69% |
| PB0168 | 1539 | 1060 | 479 | 69% |
| PB0151 | 1813 | 1251 | 562 | 69% |
| PB0022 | 776 | 537 | 239 | 69% |
| PB0225 | 714 | 496 | 218 | 70% |
| PB0251 | 415 | 289 | 126 | 70% |
| PB0122 | 964 | 672 | 292 | 70% |
| PB0157 | 418 | 293 | 125 | 70% |
| PB0188 | 224 | 157 | 67 | 70% |
| PB0150 | 811 | 571 | 240 | 70% |
| PB0018 | 1367 | 966 | 401 | 71% |
| PB0192 | 312 | 222 | 90 | 71% |
| PB0253 | 250 | 179 | 71 | 72% |
| PB0204 | 661 | 474 | 187 | 72% |
| PB0059 | 346 | 250 | 96 | 72% |
| PB0129 | 1496 | 1090 | 406 | 73% |
| PB0128 | 1792 | 1310 | 482 | 73% |

| | | | | |
|---|---|---|---|---|
| PB0201 | 413 | 303 | 110 | 73% |
| PB0202 | 249 | 183 | 66 | 74% |
| PB0036 | 485 | 362 | 123 | 75% |
| PB0005 | 383 | 287 | 96 | 75% |
| PB0065 | 1185 | 894 | 291 | 75% |
| PB0108 | 1172 | 885 | 287 | 76% |
| PB0223 | 774 | 585 | 189 | 76% |
| PB0233 | 782 | 592 | 190 | 76% |
| PB0196 | 350 | 268 | 82 | 77% |
| PB0239 | 271 | 208 | 63 | 77% |
| PB0243 | 3409 | 2642 | 767 | 78% |
| PB0006 | 303 | 235 | 68 | 78% |
| PB0130 | 574 | 447 | 127 | 78% |
| PB0173 | 993 | 777 | 216 | 78% |
| PB0236 | 826 | 646 | 180 | 78% |
| PB0161 | 2653 | 2083 | 570 | 79% |
| PB0131 | 1534 | 1229 | 305 | 80% |
| PB0160 | 442 | 354 | 88 | 80% |
| PB0078 | 364 | 293 | 71 | 81% |
| PB0237 | 627 | 519 | 108 | 83% |
| PB0205 | 333 | 276 | 57 | 83% |
| PB0019 | 1466 | 1223 | 243 | 83% |
| PB0027 | 1830 | 1543 | 287 | 84% |
| PB0011 | 335 | 287 | 48 | 86% |
| PB0248 | 1134 | 972 | 162 | 86% |
| PB0057 | 218 | 187 | 31 | 86% |
| PB0247 | 698 | 626 | 72 | 90% |
| PB0055 | 902 | 835 | 67 | 93% |
| PB0209 | 650 | 616 | 34 | 95% |
| PB0245 | 790 | 761 | 29 | 96% |
| PB0246 | 211 | 208 | 3 | 99% |

(*) Percentage of identity between *D. rerio* and *H. sapiens* protein homologs

(**) *Not found in D. rerio*

**Table S3.**
Pairwise structure alignment between *Danio rerio* and *Homo sapiens*

| Phylobone Code | Entry | Chain | RMSD | TM-score | Identity | Aligned Residues | Sequence Length | Model Residues |
|---|---|---|---|---|---|---|---|---|
| **PB0001** | AF-P02765-F1 | A | - | - | - | - | 367 | 367 |
| | AF-A5WWI5-F1 | A | 3.76 | 0.68 | 33% | 243 | 386 | 386 |
| **PB0002** | AF-P61769-F1 | A | - | - | - | - | 119 | 119 |
| | AF-B0UYS1-F1 | A | 2.07 | 0.85 | 41% | 105 | 116 | 116 |
| **PB0003** | AF-P00734-F1 | A | - | - | - | - | 622 | 622 |
| | AF-E7FAN5-F1 | A | 3.33 | 0.84 | 56% | 498 | 635 | 635 |
| **PB0004** | AF-P08493-F1 | A | - | - | - | - | 103 | 103 |
| | AF-Q6YND0-F1 | A | 4.8 | 0.35 | 5% | 48 | 105 | 105 |
| **PB0005** | AF-P21810-F1 | A | - | - | - | - | 368 | 368 |
| | AF-B7ZDB5-F1 | A | 1.86 | 0.89 | 74% | 328 | 374 | 374 |
| **PB0006** | AF-P09486-F1 | A | - | - | - | - | 303 | 303 |
| | AF-Q6IQH0-F1 | A | 1.15 | 0.78 | 84% | 237 | 291 | 291 |
| **PB0007** | AF-O15335-F1 | A | - | - | - | - | 359 | 359 |
| | AF-Q7SYE5-F1 | A | 1.76 | 0.9 | 62% | 328 | 363 | 363 |
| **PB0008** | P02743 | | | | | | | |
| | F1R6R2 | | | | | | | |
| **PB0009** | AF-P36955-F1 | A | - | - | - | - | 418 | 418 |
| | AF-Q66I20-F1 | A | 2.03 | 0.85 | 40% | 363 | 406 | 406 |
| **PB0010** | AF-P02792-F1 | A | - | - | - | - | 175 | 175 |
| | AF-Q9DDT0-F1 | A | 1.03 | 0.98 | 55% | 172 | 177 | 177 |
| **PB0011** | AF-P04406-F1 | A | - | - | - | - | 335 | 335 |
| | AF-Q5XJ10-F1 | A | 0.18 | 0.99 | 86% | 333 | 333 | 333 |
| **PB0012** | AF-P06702-F1 | A | - | - | - | - | 114 | 114 |
| | AF-Q7ZVA4-F1 | A | 1.49 | 0.78 | 38% | 96 | 100 | 100 |
| **PB0013** | AF-P08311-F1 | A | - | - | - | - | 255 | 255 |
| | AF-A8WIQ5-F1 | A | 2.03 | 0.86 | 41% | 226 | 251 | 251 |
| **PB0014** | P24158 | | | | | | | |
| | A0A8M9Q575 | | | | | | | |
| **PB0015** | AF-P28300-F1 | A | - | - | - | - | 417 | 417 |
| | AF-Q6NYT8-F1 | A | 2.72 | 0.56 | 76% | 227 | 408 | 408 |
| **PB0016** | P62805 | | | | | | | |
| | A0A8M3B0X6 | | | | | | | |
| **PB0017** | AF-P69905-F1 | A | - | - | - | - | 142 | 142 |
| | AF-Q803Z5-F1 | A | 0.81 | 0.97 | 54% | 142 | 143 | 143 |

| | | | | | | | | |
|---|---|---|---|---|---|---|---|---|
| **PB0018** | AF-P08123-F1 | A | - | - | - | - | 1366 | 1366 |
| | AF-Q6IQX2-F1 | A | 7.93 | 0.32 | 50% | 306 | 1352 | 1352 |
| **PB0019** | AF-P02452-F1 | A | - | - | - | - | 1464 | 1464 |
| | AF-F1QDL1-F1 | A | 7.41 | 0.26 | 52% | 277 | 1449 | 1449 |
| **PB0020** | AF-Q9BRR6-F1 | A | - | - | - | - | 497 | 497 |
| | AF-A0JML7-F1 | A | 1.4 | 0.89 | 67% | 444 | 498 | 498 |
| **PB0021** | AF-Q8TAL6-F1 | A | - | - | - | - | 211 | 211 |
| | AF-A1IGX5-F1 | A | 3.29 | 0.34 | 6% | 78 | 210 | 210 |
| **PB0022** | AF-Q9Y4K0-F1 | A | - | - | - | - | 774 | 774 |
| | AF-A1L1V4-F1 | A | 3.71 | 0.81 | 67% | 590 | 762 | 762 |
| **PB0023** | AF-P12645-F1 | A | - | - | - | - | 472 | 472 |
| | AF-A2AVJ4-F1 | A | 4.02 | 0.67 | 58% | 297 | 452 | 452 |
| **PB0024** | AF-O95967-F1 | A | - | - | - | - | 443 | 443 |
| | AF-A2CEN1-F1 | A | 3.62 | 0.74 | 67% | 331 | 440 | 440 |
| **PB0025** | AF-P02751-F1 | A | - | - | - | - | 2477 | 2477 |
| | AF-A2CEW3-F1 | A | 11.08 | 0.19 | 22% | 105 | 2500 | 2500 |
| **PB0026** | AF-Q08431-F1 | A | - | - | - | - | 387 | 387 |
| | AF-A2RRT1-F1 | A | 3.77 | 0.69 | 45% | 268 | 473 | 473 |
| **PB0027** | AF-P12107-F1 | A | - | - | - | - | 1806 | 1806 |
| | AF-D6MUD3-F1 | A | 8.95 | 0.22 | 38% | 252 | 1815 | 1815 |
| **PB0028** | AF-Q8N474-F1 | A | - | - | - | - | 314 | 314 |
| | AF-A3KNP6-F1 | A | 2.92 | 0.76 | 60% | 252 | 300 | 300 |
| **PB0029** | AF-Q8WXD2-F1 | A | - | - | - | - | 468 | 468 |
| | AF-A3KQQ9-F1 | A | 6.64 | 0.25 | 28% | 90 | 478 | 478 |
| **PB0030** | AF-P01024-F1 | A | - | - | - | - | 1663 | 1663 |
| | AF-B8JKW4-F1 | A | 7.61 | 0.59 | 29% | 449 | 1643 | 1643 |
| **PB0031** | AF-P21941-F1 | A | - | - | - | - | 496 | 496 |
| | AF-A5WWJ4-F1 | A | 5.77 | 0.58 | 49% | 227 | 489 | 489 |
| **PB0032** | AF-Q53GQ0-F1 | A | - | - | - | - | 312 | 312 |
| | AF-A7MCK2-F1 | A | 0.84 | 0.98 | 61% | 308 | 311 | 311 |
| **PB0033** | AF-P07355-F1 | A | - | - | - | - | 339 | 339 |
| | AF-A8E5G1-F1 | A | 1.51 | 0.91 | 66% | 318 | 338 | 338 |
| **PB0034** | AF-P36980-F1 | A | - | - | - | - | 270 | 270 |
| | AF-E2FHP6-F1 | A | 5.15 | 0.42 | 13% | 108 | 606 | 606 |
| **PB0035** | AF-P04070-F1 | A | - | - | - | - | 461 | 461 |
| | AF-A8KC28-F1 | A | 2.27 | 0.82 | 47% | 385 | 434 | 434 |
| **PB0036** | AF-Q92743-F1 | A | - | - | - | - | 480 | 480 |
| | AF-A9JRB3-F1 | A | 3.33 | 0.8 | 72% | 388 | 476 | 476 |

| | | | | | | | | |
|---|---|---|---|---|---|---|---|---|
| **PB0037** | AF-P22413-F1 | A | - | - | - | - | 925 | 925 |
| | AF-B0JZL7-F1 | A | 3.18 | 0.86 | 53% | 762 | 878 | 878 |
| **PB0038** | AF-P24593-F1 | A | - | - | - | - | 272 | 272 |
| | AF-B0S525-F1 | A | 3.63 | 0.73 | 52% | 206 | 265 | 265 |
| **PB0039** | AF-O75829-F1 | A | - | - | - | - | 334 | 334 |
| | AF-P58239-F1 | A | 5.12 | 0.34 | 29% | 108 | 286 | 286 |
| **PB0040** | AF-P27658-F1 | A | - | - | - | - | 744 | 744 |
| | AF-B7SW26-F1 | A | 6.62 | 0.27 | 56% | 151 | 711 | 711 |
| **PB0041** | AF-Q99645-F1 | A | - | - | - | - | 322 | 322 |
| | AF-B0S5X0-F1 | A | 2.57 | 0.67 | 60% | 210 | 712 | 712 |
| **PB0042** | AF-P48307-F1 | A | - | - | - | - | 235 | 235 |
| | AF-B0S6C1-F1 | A | 5.33 | 0.29 | 26% | 67 | 279 | 279 |
| **PB0043** | AF-Q9UQP3-F1 | A | - | - | - | - | 1299 | 1299 |
| | AF-B0S6K5-F1 | A | 5.76 | 0.35 | 45% | 306 | 932 | 932 |
| **PB0044** | AF-P29400-F1 | A | - | - | - | - | 1685 | 1685 |
| | AF-B0UXF7-F1 | A | 8.11 | 0.23 | 48% | 247 | 1679 | 1679 |
| **PB0045** | AF-O00391-F1 | A | - | - | - | - | 747 | 747 |
| | AF-B0UXN0-F1 | A | 2.67 | 0.67 | 48% | 495 | 778 | 778 |
| **PB0046** | AF-Q68BL7-F1 | A | - | - | - | - | 652 | 652 |
| | AF-B0UXR7-F1 | A | 5.12 | 0.56 | 59% | 314 | 645 | 645 |
| **PB0047** | AF-P02652-F1 | A | - | - | - | - | 100 | 100 |
| | AF-B3DFP9-F1 | A | 4.37 | 0.41 | 11% | 50 | 141 | 141 |
| **PB0048** | AF-O43405-F1 | A | - | - | - | - | 550 | 550 |
| | AF-B3DFV8-F1 | A | 1.13 | 0.7 | 64% | 389 | 553 | 553 |
| **PB0049** | AF-P06727-F1 | A | - | - | - | - | 396 | 396 |
| | AF-B3DHC5-F1 | A | 4.62 | 0.31 | 14% | 121 | 255 | 255 |
| **PB0050** | AF-Q15113-F1 | A | - | - | - | - | 449 | 449 |
| | AF-B3DJG2-F1 | A | 4.95 | 0.51 | 43% | 219 | 538 | 538 |
| **PB0051** | Q99715 | | | | | | | |
| | E7FG81 | | | | | | | |
| **PB0052** | AF-Q76M96-F1 | A | - | - | - | - | 950 | 950 |
| | AF-B7SDQ7-F1 | A | 6.97 | 0.26 | 39% | 181 | 867 | 867 |
| **PB0053** | AF-P55268-F1 | A | - | - | - | - | 1798 | 1798 |
| | AF-B7ZDA6-F1 | A | 5.72 | 0.25 | 61% | 399 | 1782 | 1782 |
| **PB0054** | AF-P20849-F1 | A | - | - | - | - | 921 | 921 |
| | AF-F1QTQ2-F1 | A | 8.24 | 0.1 | 9% | 48 | 687 | 687 |
| **PB0055** | AF-P12814-F1 | A | - | - | - | - | 892 | 892 |
| | AF-B8JHU4-F1 | A | 8.24 | 0.97 | 93% | 863 | 902 | 902 |

| | | | | | | | | |
|---|---|---|---|---|---|---|---|---|
| **PB0056** | AF-P08294-F1 | A | - | - | - | - | 240 | 240 |
| | AF-E9QIM2-F1 | A | 1.62 | 0.66 | 52% | 158 | 192 | 192 |
| **PB0057** | AF-P26583-F1 | A | - | - | - | - | 209 | 209 |
| | AF-B8JL29-F1 | A | 4 | 0.48 | 58% | 97 | 213 | 213 |
| **PB0058** | AF-P02787-F1 | A | - | - | - | - | 698 | 698 |
| | AF-B8JL43-F1 | A | 3.79 | 0.82 | 40% | 525 | 673 | 673 |
| **PB0059** | AF-P04083-F1 | A | - | - | - | - | 346 | 346 |
| | AF-B8JLZ3-F1 | A | 2.52 | 0.92 | 52% | 314 | 342 | 342 |
| **PB0060** | AF-O00584-F1 | A | - | - | - | - | 256 | 256 |
| | AF-B8XY56-F1 | A | 2.18 | 0.77 | 48% | 200 | 240 | 240 |
| **PB0061** | AF-Q8WWA0-F1 | A | - | - | - | - | 313 | 313 |
| | AF-C1IHU8-F1 | A | 1.22 | 0.86 | 50% | 274 | 311 | 311 |
| **PB0062** | AF-B2RNN3-F1 | A | - | - | - | - | 333 | 333 |
| | AF-E7F7K2-F1 | A | 3.58 | 0.46 | 52% | 148 | 339 | 339 |
| **PB0063** | P22105 | | | | | | | |
| | E7EXE8 | | | | | | | |
| **PB0064** | AF-P05452-F1 | A | - | - | - | - | 202 | 202 |
| | AF-E7EYJ8-F1 | A | 1.61 | 0.84 | 55% | 178 | 197 | 197 |
| **PB0065** | AF-P35442-F1 | A | - | - | - | - | 1172 | 1172 |
| | AF-E7F1S4-F1 | A | 4.62 | 0.75 | 65% | 758 | 1181 | 1181 |
| **PB0066** | AF-P08582-F1 | A | - | - | - | - | 738 | 738 |
| | AF-E7F2E3-F1 | A | 4.64 | 0.76 | 44% | 513 | 723 | 723 |
| **PB0067** | AF-P00740-F1 | A | - | - | - | - | 461 | 461 |
| | AF-E7F301-F1 | A | 2.93 | 0.65 | 43% | 301 | 503 | 503 |
| **PB0068** | AF-P16112-F1 | A | - | - | - | - | 2530 | 2530 |
| | AF-F1R511-F1 | A | 6.28 | 0.19 | 47% | 345 | 1207 | 1207 |
| **PB0069** | AF-Q15063-F1 | A | - | - | - | - | 836 | 836 |
| | AF-Q75U66-F1 | A | 3.44 | 0.74 | 61% | 628 | 782 | 782 |
| **PB0070** | AF-Q14112-F1 | A | - | - | - | - | 1375 | 1375 |
| | AF-X1WE42-F1 | A | 7.34 | 0.39 | 32% | 259 | 1576 | 1576 |
| **PB0071** | AF-Q9UMD9-F1 | A | - | - | - | - | 1497 | 1497 |
| | AF-B8A4S4-F1 | A | 10.74 | 0.12 | 14% | 59 | 1408 | 1408 |
| **PB0072** | AF-Q8IUX7-F1 | A | - | - | - | - | 1158 | 1158 |
| | AF-E7F829-F1 | A | 3.1 | 0.56 | 59% | 609 | 1022 | 1022 |
| **PB0073** | AF-Q8NHW6-F1 | A | - | - | - | - | 89 | 89 |
| | AF-A0A286Y9F3-F1 | A | 2.11 | 0.57 | 55% | 55 | 90 | 90 |
| **PB0074** | AF-O60687-F1 | A | - | - | - | - | 465 | 465 |
| | AF-E7F8X0-F1 | A | 2.72 | 0.85 | 56% | 417 | 460 | 460 |

| | | | | | | | | |
|---|---|---|---|---|---|---|---|---|
| **PB0075** | AF-P14543-F1 | A | - | - | - | - | 1247 | 1247 |
| | AF-F1RAG3-F1 | A | 7.14 | 0.44 | 31% | 377 | 1175 | 1175 |
| **PB0076** | AF-Q96P44-F1 | A | - | - | - | - | 957 | 957 |
| | AF-E7FB76-F1 | A | 3.77 | 0.47 | 57% | 423 | 717 | 717 |
| **PB0077** | P04114 | | | | | | | |
| | Q5TZ29 | | | | | | | |
| **PB0078** | AF-P10915-F1 | A | - | - | - | - | 354 | 354 |
| | AF-E7FBW8-F1 | A | 2.05 | 0.91 | 75% | 320 | 353 | 353 |
| **PB0079** | P25940 | | | | | | | |
| | D6MUD7 | | | | | | | |
| **PB0080** | AF-P01031-F1 | A | - | - | - | - | 1676 | 1676 |
| | AF-E7FCV2-F1 | A | 5.35 | 0.77 | 33% | 989 | 1704 | 1704 |
| **PB0081** | AF-P12110-F1 | A | - | - | - | - | 1019 | 1019 |
| | AF-E7FCV8-F1 | A | 8.3 | 0.34 | 17% | 136 | 1015 | 1015 |
| **PB0082** | AF-Q7Z5L7-F1 | A | - | - | - | - | 613 | 613 |
| | AF-E7FDK4-F1 | A | 2.74 | 0.76 | 60% | 462 | 552 | 552 |
| **PB0083** | Q9HC84 | | | | | | | |
| | X1WG18 | | | | | | | |
| **PB0084** | AF-P05997-F1 | A | - | - | - | - | 1499 | 1499 |
| | AF-E7FE47-F1 | A | 8.42 | 0.26 | 33% | 262 | 1499 | 1499 |
| **PB0085** | AF-Q9UGM5-F1 | A | - | - | - | - | 382 | 382 |
| | AF-E7FE90-F1 | A | 3.63 | 0.78 | 32% | 293 | 499 | 499 |
| **PB0086** | AF-P02647-F1 | A | - | - | - | - | 267 | 267 |
| | AF-A0A0R4IKF0-F1 | A | 4.07 | 0.39 | 13% | 104 | 257 | 257 |
| **PB0087** | AF-P02654-F1 | A | - | - | - | - | 83 | 83 |
| | AF-E9QDI1-F1 | A | 2.87 | 0.6 | 31% | 59 | 85 | 85 |
| **PB0088** | AF-P02655-F1 | A | - | - | - | - | 101 | 101 |
| | AF-E9QEQ1-F1 | A | 4.14 | 0.43 | 8% | 58 | 100 | 100 |
| **PB0089** | AF-Q13103-F1 | A | - | - | - | - | 211 | 211 |
| | AF-E9QG22-F1 | A | 2.9 | 0.49 | 27% | 110 | 194 | 194 |
| **PB0090** | B7ZW00 | | | | | | | |
| | F1Q4X1 | | | | | | | |
| **PB0091** | Q07092 | | | | | | | |
| | A0A8M9PNN1 | | | | | | | |
| **PB0092** | AF-P05186-F1 | A | - | - | - | - | 524 | 524 |
| | AF-F1Q5B5-F1 | A | 1.41 | 0.94 | 75% | 491 | 561 | 561 |
| **PB0093** | Q6UXX5 | | | | | | | |
| | A0A8M3AYJ4 | | | | | | | |

| | | | | | | | | |
|---|---|---|---|---|---|---|---|---|
| **PB0094** | AF-Q8IUL8-F1 | A | - | - | - | - | 1156 | 1156 |
| | AF-F1Q5N7-F1 | A | 3.6 | 0.85 | 55% | 933 | 1255 | 1255 |
| **PB0095** | AF-P22891-F1 | A | - | - | - | - | 400 | 400 |
| | AF-F1Q5S2-F1 | A | 3.01 | 0.67 | 30% | 266 | 426 | 426 |
| **PB0096** | AF-P12109-F1 | A | - | - | - | - | 1028 | 1028 |
| | AF-F1Q6P3-F1 | A | 5.44 | 0.44 | 47% | 345 | 1007 | 1007 |
| **PB0097** | AF-Q8N2E2-F1 | A | - | - | - | - | 1590 | 1590 |
| | AF-A0A0R4IWA2-F1 | A | 4.89 | 0.55 | 46% | 774 | 1721 | 1721 |
| **PB0098** | AF-Q06828-F1 | A | - | - | - | - | 376 | 376 |
| | AF-F1QG51-F1 | A | 1.48 | 0.78 | 56% | 303 | 342 | 342 |
| **PB0099** | AF-O75339-F1 | A | - | - | - | - | 1184 | 1184 |
| | AF-F1Q775-F1 | A | 4.4 | 0.82 | 62% | 901 | 1171 | 1171 |
| **PB0100** | AF-P43235-F1 | A | - | - | - | - | 329 | 329 |
| | AF-F1Q8A0-F1 | A | 1.19 | 0.93 | 64% | 309 | 333 | 333 |
| **PB0101** | AF-A6NMZ7-F1 | A | - | - | - | - | 2263 | 2263 |
| | AF-X1WEZ4-F1 | A | 5.82 | 0.44 | 34% | 798 | 1756 | 1756 |
| **PB0102** | AF-P10451-F1 | A | - | - | - | - | 314 | 314 |
| | AF-Q6IVB9-F1 | A | 7.23 | 0.15 | 0.15 | 33 | 305 | 305 |
| **PB0103** | AF-Q641Q3-F1 | A | - | - | - | - | 311 | 311 |
| | AF-Q7ZV46-F1 | A | 2.87 | 0.77 | 56% | 251 | 286 | 286 |
| **PB0104** | AF-Q96A84-F1 | A | - | - | - | - | 441 | 441 |
| | AF-A0A286Y830-F1 | A | 6.44 | 0.21 | 6% | 75 | 354 | 354 |
| **PB0105** | AF-A4D0S4-F1 | A | - | - | - | - | 1761 | 1761 |
| | AF-Q8JHV6-F1 | A | 6.48 | 0.28 | 50% | 398 | 1827 | 1827 |
| **PB0106** | AF-O15444-F1 | A | - | - | - | - | 150 | 150 |
| | AF-F1QBE9-F1 | A | 2.33 | 0.45 | 37% | 73 | 106 | 106 |
| **PB0107** | AF-P14780-F1 | A | - | - | - | - | 707 | 707 |
| | AF-F1QC76-F1 | A | 3.36 | 0.59 | 62% | 402 | 680 | 680 |
| **PB0108** | AF-F1QC76-F1 | A | - | - | - | - | 680 | 680 |
| | AF-F1QEE7-F1 | A | 8.5 | 0.24 | 4% | 70 | 1170 | 1170 |
| **PB0109** | O95445 | | | | | | | |
| | F1QFH8 | | | | | | | |
| **PB0110** | AF-P08709-F1 | A | - | - | - | - | 466 | 466 |
| | AF-F1QFP3-F1 | A | 2.63 | 0.65 | 44% | 304 | 433 | 433 |
| **PB0111** | AF-Q9NS15-F1 | A | - | - | - | - | 1303 | 1303 |
| | AF-F1QFX6-F1 | A | 7.48 | 0.2 | 42% | 181 | 1258 | 1258 |
| **PB0112** | AF-P45452-F1 | A | - | - | - | - | 471 | 471 |
| | AF-Q71G59-F1 | A | 2.04 | 0.88 | 54% | 418 | 475 | 475 |

| | | | | | | | | |
|---|---|---|---|---|---|---|---|---|
| **PB0113** | AF-P02462-F1 | A | - | - | - | - | 1669 | 1669 |
| | AF-F1QG79-F1 | A | 8.81 | 0.25 | 51% | 182 | 1317 | 1317 |
| **PB0114** | AF-Q14393-F1 | A | - | - | - | - | 678 | 678 |
| | AF-F1QGG9-F1 | A | 4.3 | 0.71 | 46% | 454 | 648 | 648 |
| **PB0115** | AF-Q15198-F1 | A | - | - | - | - | 375 | 375 |
| | AF-F1QHI8-F1 | A | 2.86 | 0.57 | 54% | 213 | 371 | 371 |
| **PB0116** | P04004 | | | | | | | |
| | A0A8M9QLE5 | | | | | | | |
| **PB0117** | AF-Q13753-F1 | A | - | - | - | - | 1193 | 1193 |
| | AF-F1QIJ3-F1 | A | 8.82 | 0.19 | 6% | 72 | 736 | 736 |
| **PB0118** | AF-P06681-F1 | A | - | - | - | - | 752 | 752 |
| | AF-F1QJB3-F1 | A | 5.81 | 0.69 | 30% | 414 | 831 | 831 |
| **PB0119** | AF-Q5T4F7-F1 | A | - | - | - | - | 317 | 317 |
| | AF-F1QJG6-F1 | A | 3.69 | 0.65 | 57% | 225 | 310 | 310 |
| **PB0120** | AF-P07225-F1 | A | - | - | - | - | 676 | 676 |
| | AF-F1QK29-F1 | A | 4.36 | 0.8 | 46% | 495 | 673 | 673 |
| **PB0121** | AF-O75443-F1 | A | - | - | - | - | 2155 | 2155 |
| | AF-F1QK57-F1 | A | 5.71 | 0.36 | 55% | 693 | 2201 | 2201 |
| **PB0122** | AF-P49746-F1 | A | - | - | - | - | 956 | 956 |
| | AF-F1QKL1-F1 | A | 4.74 | 0.62 | 69% | 538 | 962 | 962 |
| **PB0123** | O15230 | | | | | | | |
| | F1QKW3 | | | | | | | |
| **PB0124** | AF-P00742-F1 | A | - | - | - | - | 488 | 488 |
| | AF-F1QLC3-F1 | A | 4.32 | 0.67 | 46% | 304 | 504 | 504 |
| **PB0125** | AF-Q02818-F1 | A | - | - | - | - | 461 | 461 |
| | AF-F1QLH9-F1 | A | 3.53 | 0.76 | 65% | 351 | 454 | 454 |
| **PB0126** | AF-Q9BYJ0-F1 | A | - | - | - | - | 223 | 223 |
| | AF-F1QLT3-F1 | A | 2.98 | 0.51 | 32% | 117 | 279 | 279 |
| **PB0127** | AF-P20774-F1 | A | - | - | - | - | 298 | 298 |
| | AF-F1QLW6-F1 | A | 1.16 | 0.71 | 61% | 211 | 245 | 245 |
| **PB0128** | P07942 | | | | | | | |
| | F1REL9 | | | | | | | |
| **PB0129** | AF-P02458-F1 | A | - | - | - | - | 1487 | 1487 |
| | AF-Q2LDA1-F1 | A | 8.46 | 0.26 | 43% | 261 | 1491 | 1491 |
| **PB0130** | AF-Q8TB73-F1 | A | - | - | - | - | 568 | 568 |
| | AF-F1QPX0-F1 | A | 2.27 | 0.93 | 78% | 532 | 574 | 574 |
| **PB0131** | O94813 | | | | | | | |
| | F1QQE4 | | | | | | | |

| | | | | | | | | |
|---|---|---|---|---|---|---|---|---|
| **PB0132** | AF-P05121-F1 | A | - | - | - | - | 402 | 402 |
| | AF-F1QRB8-F1 | A | 1.58 | 0.9 | 44% | 368 | 392 | 392 |
| **PB0133** | AF-P55001-F1 | A | - | - | - | - | 183 | 183 |
| | AF-B3DKC3-F1 | A | 3.6 | 0.35 | 73% | 69 | 158 | 158 |
| **PB0134** | AF-Q5GFL6-F1 | A | - | - | - | - | 755 | 755 |
| | AF-U3JA51-F1 | A | 5.93 | 0.7 | 40% | 382 | 795 | 795 |
| **PB0135** | Q16787 | | | | | | | |
| | A0A2R8Q7H6 | | | | | | | |
| **PB0136** | AF-Q14520-F1 | A | - | - | - | - | 560 | 560 |
| | AF-F1QW52-F1 | A | 4.78 | 0.67 | 42% | 339 | 546 | 546 |
| **PB0137** | AF-Q03692-F1 | A | - | - | - | - | 680 | 680 |
| | AF-F1QXD5-F1 | A | 6.19 | 0.27 | 36% | 150 | 655 | 655 |
| **PB0138** | AF-P51888-F1 | A | - | - | - | - | 382 | 382 |
| | AF-F1QY29-F1 | A | 1.72 | 0.86 | 60% | 332 | 378 | 378 |
| **PB0139** | AF-P24821-F1 | A | - | - | - | - | 2201 | 2201 |
| | AF-F1QYE2-F1 | A | 7.16 | 0.23 | 50% | 376 | 1811 | 1811 |
| **PB0140** | AF-Q14031-F1 | A | - | - | - | - | 1691 | 1691 |
| | AF-F1QYN1-F1 | A | 9.05 | 0.26 | 44% | 202 | 1714 | 1714 |
| **PB0141** | AF-Q16363-F1 | A | - | - | - | - | 1823 | 1823 |
| | AF-F1QYU0-F1 | A | 6.74 | 0.45 | 45% | 569 | 1871 | 1871 |
| **PB0142** | AF-O00468-F1 | A | - | - | - | - | 2068 | 2068 |
| | AF-F1R074-F1 | A | 10.15 | 0.27 | 26% | 258 | 2026 | 2026 |
| **PB0143** | AF-Q8N6Y2-F1 | A | - | - | - | - | 441 | 441 |
| | AF-F1R0N0-F1 | A | 3.97 | 0.44 | 51% | 180 | 435 | 435 |
| **PB0144** | AF-Q9BX93-F1 | A | - | - | - | - | 195 | 195 |
| | AF-F1R147-F1 | A | 2.11 | 0.81 | 65% | 164 | 235 | 235 |
| **PB0145** | AF-Q14050-F1 | A | - | - | - | - | 684 | 684 |
| | AF-F1R1F1-F1 | A | 8.71 | 0.26 | 16% | 74 | 679 | 679 |
| **PB0146** | AF-P35443-F1 | A | - | - | - | - | 961 | 961 |
| | AF-F1R1P9-F1 | A | 3.22 | 0.58 | 82% | 535 | 949 | 949 |
| **PB0147** | AF-P08572-F1 | A | - | - | - | - | 1712 | 1712 |
| | AF-A0A0H2UKP2-F1 | A | 8.67 | 0.23 | 44% | 255 | 1669 | 1669 |
| **PB0148** | AF-O94769-F1 | A | - | - | - | - | 699 | 699 |
| | AF-F1R2S9-F1 | A | 3.29 | 0.58 | 59% | 394 | 731 | 731 |
| **PB0149** | AF-Q6UX07-F1 | A | - | - | - | - | 377 | 377 |
| | AF-F1R2W1-F1 | A | 1.6 | 0.8 | 54% | 306 | 318 | 318 |

| | | | | | | | | |
|---|---|---|---|---|---|---|---|---|
| **PB0150** | AF-Q9HCB6-F1 | A | - | - | - | - | 807 | 807 |
| | AF-F1R2Z9-F1 | A | 5.93 | 0.54 | 46% | 328 | 804 | 804 |
| **PB0151** | P13942 | | | | | | | |
| | D6MUD4 | | | | | | | |
| **PB0152** | P98160 | | | | | | | |
| | F1R5K7 | | | | | | | |
| **PB0153** | AF-Q01955-F1 | A | - | - | - | - | 1670 | 1670 |
| | AF-F1R5V4-F1 | A | 9.42 | 0.12 | 18% | 57 | 920 | 920 |
| **PB0154** | AF-Q4ZHG4-F1 | A | - | - | - | - | 1894 | 1894 |
| | AF-F1R5Z2-F1 | A | 7.04 | 0.22 | 42% | 318 | 1645 | 1645 |
| **PB0155** | AF-P51884-F1 | A | - | - | - | - | 338 | 338 |
| | AF-Q6IQQ7-F1 | A | 2.03 | 0.91 | 50% | 312 | 344 | 344 |
| **PB0156** | P35555 | | | | | | | |
| | A0A8M6YW17 | | | | | | | |
| **PB0157** | AF-Q9UKZ9-F1 | A | - | - | - | - | 415 | 415 |
| | AF-F1R774-F1 | A | 5.28 | 0.68 | 50% | 246 | 413 | 413 |
| **PB0158** | AF-O95841-F1 | A | - | - | - | - | 491 | 491 |
| | AF-Q5KQU0-F1 | A | 2.78 | 0.47 | 69% | 225 | 480 | 480 |
| **PB0159** | AF-P00751-F1 | A | - | - | - | - | 764 | 764 |
| | AF-F1R886-F1 | A | 5.5 | 0.72 | 35% | 426 | 761 | 761 |
| **PB0160** | AF-P61812-F1 | A | - | - | - | - | 414 | 414 |
| | AF-Q7SZV4-F1 | A | 2.92 | 0.85 | 77% | 344 | 411 | 411 |
| **PB0161** | AF-P21333-F1 | A | - | - | - | - | 2647 | 2647 |
| | AF-E9QI62-F1 | A | 11.06 | 0.29 | 44% | 166 | 2553 | 2553 |
| **PB0162** | AF-P05156-F1 | A | - | - | - | - | 583 | 583 |
| | AF-A0A0G2L5F9-F1 | A | 2.74 | 0.66 | 45% | 377 | 428 | 428 |
| **PB0163** | AF-P21333-F1 | A | - | - | - | - | 2647 | 2647 |
| | AF-E9QI62-F1 | A | 11.06 | 0.29 | 44% | 166 | 2553 | 2553 |
| **PB0164** | AF-Q05707-F1 | A | - | - | - | - | 1796 | 1796 |
| | AF-F1R9Y1-F1 | A | 3.72 | 0.32 | 39% | 224 | 1864 | 1864 |
| **PB0165** | AF-P08697-F1 | A | - | - | - | - | 491 | 491 |
| | AF-F1RAP6-F1 | A | 2.78 | 0.71 | 40% | 351 | 480 | 480 |
| **PB0166** | AF-O94907-F1 | A | - | - | - | - | 266 | 266 |

| | | | | | | | | |
|---|---|---|---|---|---|---|---|---|
| | AF-Q9PWH3-F1 | A | 4.57 | 0.47 | 58% | 125 | 240 | 240 |
| **PB0167** | AF-Q66K79-F1 | A | - | - | - | - | 652 | 652 |
| | AF-F6P711-F1 | A | 4.61 | 0.75 | 49% | 443 | 653 | 653 |
| **PB0168** | AF-Q66K79-F1 | A | - | - | - | - | 652 | 652 |
| | AF-F6P711-F1 | A | 4.61 | 0.75 | 49% | 443 | 653 | 653 |
| **PB0169** | Q9Y6R7 | | | | | | | |
| | Q9Y6R7 | | | | | | | |
| **PB0170** | Q75N90 | | | | | | | |
| | A0A0G2KQ62 | | | | | | | |
| **PB0171** | AF-P83110-F1 | A | - | - | - | - | 453 | 453 |
| | AF-F1REI0-F1 | A | 3.03 | 0.46 | 72% | 200 | 416 | 416 |
| **PB0172** | AF-Q9NQ30-F1 | A | - | - | - | - | 184 | 184 |
| | AF-A0A0R4IF38-F1 | A | 2.58 | 0.63 | 54% | 124 | 179 | 179 |
| **PB0173** | AF-P13497-F1 | A | - | - | - | - | 986 | 986 |
| | AF-F1REM3-F1 | A | 4.73 | 0.54 | 73% | 495 | 976 | 976 |
| **PB0174** | AF-P07477-F1 | A | - | - | - | - | 247 | 247 |
| | AF-Q8AV83-F1 | A | 2.09 | 0.9 | 68% | 221 | 243 | 243 |
| **PB0175** | AF-P01009-F1 | A | - | - | - | - | 418 | 418 |
| | AF-Q5SPJ4-F1 | A | 1.83 | 0.86 | 42% | 367 | 429 | 429 |
| **PB0176** | AF-Q15582-F1 | A | - | - | - | - | 683 | 683 |
| | AF-Q503K1-F1 | A | 2.99 | 0.86 | 64% | 594 | 677 | 677 |
| **PB0177** | AF-P20062-F1 | A | - | - | - | - | 427 | 427 |
| | AF-F2Z4S3-F1 | A | 2.64 | 0.84 | 31% | 361 | 423 | 423 |
| **PB0178** | AF-O14793-F1 | A | - | - | - | - | 375 | 375 |
| | AF-O42222-F1 | A | 3.37 | 0.48 | 72% | 177 | 374 | 374 |
| **PB0179** | AF-P02649-F1 | A | - | - | - | - | 317 | 317 |
| | AF-O42364-F1 | A | 4.36 | 0.46 | 19% | 154 | 281 | 281 |
| **PB0180** | AF-Q9UBM4-F1 | A | - | - | - | - | 332 | 332 |
| | AF-Q15JE7-F1 | A | 2.29 | 0.63 | 53% | 209 | 322 | 322 |
| **PB0181** | AF-Q9GZV9-F1 | A | - | - | - | - | 251 | 251 |
| | AF-Q1L8M3-F1 | A | 3.56 | 0.6 | 31% | 163 | 258 | 258 |
| **PB0182** | AF-O75718-F1 | A | - | - | - | - | 401 | 401 |

| | | | | | | | | |
|---|---|---|---|---|---|---|---|---|
| | AF-Q1L8P0-F1 | A | 1.74 | 0.91 | 68% | 362 | 396 | 396 |
| **PB0183** | AF-O75718-F1 | A | - | - | - | - | 401 | 401 |
| | AF-Q1L8P0-F1 | A | 1.74 | 0.91 | 68% | 362 | 396 | 396 |
| **PB0184** | AF-P11047-F1 | A | - | - | - | - | 1609 | 1609 |
| | AF-Q1LVF0-F1 | A | 9.93 | 0.38 | 51% | 172 | 1593 | 1593 |
| **PB0185** | AF-Q9BXN1-F1 | A | - | - | - | - | 380 | 380 |
| | AF-Q1LXA7-F1 | A | 2.5 | 0.85 | 55% | 318 | 370 | 370 |
| **PB0186** | AF-Q14055-F1 | A | - | - | - | - | 689 | 689 |
| | AF-Q1LXU2-F1 | A | 8.81 | 0.14 | 9% | 39 | 691 | 691 |
| **PB0187** | AF-P01042-F1 | A | - | - | - | - | 644 | 644 |
| | AF-Q1LYJ7-F1 | A | 3.89 | 0.19 | 26% | 113 | 331 | 331 |
| **PB0188** | AF-P16035-F1 | A | - | - | - | - | 220 | 220 |
| | AF-Q1RLX2-F1 | A | 1.47 | 0.95 | 71% | 215 | 220 | 220 |
| **PB0189** | AF-P61626-F1 | A | - | - | - | - | 148 | 148 |
| | AF-Q24JW2-F1 | A | 1.72 | 0.83 | 40% | 125 | 151 | 151 |
| **PB0190** | AF-Q9NRN5-F1 | A | - | - | - | - | 406 | 406 |
| | AF-Q29RB4-F1 | A | 2.22 | 0.65 | 51% | 265 | 390 | 390 |
| **PB0191** | AF-Q07507-F1 | A | - | - | - | - | 201 | 201 |
| | AF-Q29RF0-F1 | A | 1.41 | 0.79 | 65% | 161 | 183 | 183 |
| **PB0192** | AF-Q12841-F1 | A | - | - | - | - | 308 | 308 |
| | AF-Q2PMI2-F1 | A | 1.84 | 0.54 | 72% | 165 | 310 | 310 |
| **PB0193** | AF-Q99972-F1 | A | - | - | - | - | 504 | 504 |
| | AF-Q5F0G5-F1 | A | 3.59 | 0.56 | 52% | 271 | 474 | 474 |
| **PB0194** | AF-O95460-F1 | A | - | - | - | - | 622 | 622 |
| | AF-Q5NJJ5-F1 | A | 4.41 | 0.37 | 52% | 210 | 944 | 944 |
| **PB0195** | AF-O15232-F1 | A | - | - | - | - | 486 | 486 |
| | AF-Q68EK6-F1 | A | 2.82 | 0.46 | 61% | 217 | 337 | 337 |
| **PB0196** | AF-P29279-F1 | A | - | - | - | - | 349 | 349 |
| | AF-Q5RI33-F1 | A | 4.48 | 0.52 | 72% | 171 | 345 | 345 |
| **PB0197** | AF-O60938-F1 | A | - | - | - | - | 352 | 352 |
| | AF-Q5RI43-F1 | A | 1.64 | 0.88 | 59% | 314 | 348 | 348 |
| **PB0198** | AF-P07585-F1 | A | - | - | - | - | 359 | 359 |
| | AF-Q5RI45-F1 | A | 1.91 | 0.87 | 69% | 310 | 373 | 373 |

| | | | | | | | | |
|---|---|---|---|---|---|---|---|---|
| **PB0199** | AF-Q9UK55-F1 | A | - | - | - | - | 444 | 444 |
| | AF-Q5RIH8-F1 | A | 1.85 | 0.81 | 29% | 364 | 391 | 391 |
| **PB0200** | AF-P10909-F1 | A | - | - | - | - | 449 | 449 |
| | AF-Q5SPR2-F1 | A | 5.48 | 0.62 | 30% | 235 | 449 | 449 |
| **PB0201** | AF-P10600-F1 | A | - | - | - | - | 412 | 412 |
| | AF-Q66I23-F1 | A | 2.36 | 0.9 | 74% | 369 | 410 | 410 |
| **PB0202** | AF-Q96CG8-F1 | A | - | - | - | - | 243 | 243 |
| | AF-Q96CG8-F1 | A | 0 | 1 | 100% | 243 | 243 | 243 |
| **PB0203** | AF-Q9UNI1-F1 | A | - | - | - | - | 258 | 258 |
| | AF-Q6AZC0-F1 | A | 1.87 | 0.94 | 63% | 240 | 266 | 266 |
| **PB0204** | AF-P08253-F1 | A | - | - | - | - | 660 | 660 |
| | AF-Q6DG10-F1 | A | 2.54 | 0.93 | 71% | 593 | 657 | 657 |
| **PB0205** | AF-O43852-F1 | A | - | - | - | - | 315 | 315 |
| | AF-Q6IQP3-F1 | A | 4.05 | 0.68 | 77% | 224 | 315 | 315 |
| **PB0206** | AF-Q8WVF2-F1 | A | - | - | - | - | 138 | 138 |
| | AF-Q6NWB6-F1 | A | 3.32 | 0.61 | 51% | 94 | 135 | |
| **PB0207** | AF-Q15165-F1 | A | - | - | - | - | 354 | 354 |
| | AF-Q6NXA5-F1 | A | 0.89 | 0.98 | 53% | 352 | 355 | 355 |
| **PB0208** | AF-P02675-F1 | A | - | - | - | - | 491 | 491 |
| | AF-Q6NYE1-F1 | A | 2.6 | 0.8 | 55% | 386 | 485 | 485 |
| **PB0209** | AF-P11142-F1 | A | - | - | - | - | 646 | 646 |
| | AF-Q6NYR4-F1 | A | 1.21 | 0.96 | 96% | 617 | 649 | 649 |
| **PB0210** | No danio | | | | | | | |
| **PB0211** | AF-P08758-F1 | A | - | - | - | - | 320 | 320 |
| | AF-Q6P0V8-F1 | A | 0.74 | 0.98 | 64% | 316 | 317 | 317 |
| **PB0212** | AF-P02753-F1 | A | - | - | - | - | 201 | 201 |
| | AF-Q6PC07-F1 | A | 1.48 | 0.86 | 52% | 176 | 196 | 196 |
| **PB0213** | AF-O60565-F1 | A | - | - | - | - | 184 | 184 |

| | | | | | | | | |
|---|---|---|---|---|---|---|---|---|
| | AF-Q6T937-F1 | A | 2.38 | 0.54 | 66% | 101 | 177 | 177 |
| **PB0214** | AF-P02679-F1 | A | - | - | - | - | 453 | 453 |
| | AF-Q7ZVG7-F1 | A | 2.7 | 0.86 | 56% | 384 | 431 | 431 |
| **PB0215** | AF-P07093-F1 | A | - | - | - | - | 398 | 398 |
| | AF-Q7ZVL5-F1 | A | 1.53 | 0.91 | 54% | 367 | 395 | 395 |
| **PB0216** | AF-Q9Y287-F1 | A | - | - | - | - | 266 | 266 |
| | AF-Q803H7-F1 | A | 2.52 | 0.73 | 54% | 197 | 261 | 261 |
| **PB0217** | AF-Q6UWW0-F1 | A | - | - | - | - | 184 | 184 |
| | AF-Q8QGV5-F1 | A | 2.47 | 0.83 | 26% | 160 | 184 | 184 |
| **PB0218** | AF-P21741-F1 | A | - | - | - | - | 143 | 143 |
| | AF-Q9W767-F1 | A | 3.95 | 0.4 | 68% | 54 | 146 | 146 |
| **PB0219** | AF-P18065-F1 | A | - | - | - | - | 325 | 325 |
| | AF-Q9PTH3-F1 | A | 4.83 | 0.37 | 42% | 111 | 276 | 276 |
| **PB0220** | AF-Q92765-F1 | A | - | - | - | - | 325 | 325 |
| | AF-Q9W6E0-F1 | A | 4.15 | 0.5 | 44% | 152 | 315 | 315 |
| **PB0221** | AF-P19883-F1 | A | - | - | - | - | 344 | 344 |
| | AF-Q9YHV4-F1 | A | 1.41 | 0.88 | 77% | 307 | 322 | 322 |
| **PB0222** | P01023 | | | | | | | |
| | B8QSI4 | | | | | | | |
| **PB0223** | AF-P05067-F1 | A | - | - | - | - | 770 | 770 |
| | AF-I6ZM20-F1 | A | 6.26 | 0.39 | 51% | 241 | 682 | 682 |
| **PB0224** | AF-P40121-F1 | A | - | - | - | - | 348 | 348 |
| | AF-A0A2R8Q5S0-F1 | A | 2.9 | 0.87 | 54% | 310 | 362 | 362 |
| **PB0225** | AF-P07384-F1 | A | - | - | - | - | 714 | 714 |
| | AF-Q7ZUR1-F1 | A | 0.84 | 0.98 | 71% | 701 | 704 | 704 |
| **PB0226** | AF-P17655-F1 | A | - | - | - | - | 700 | 700 |
| | AF-Q5BLH5-F1 | A | 0.85 | 0.99 | 67% | 698 | 698 | 698 |
| **PB0227** | AF-P04632-F1 | A | - | - | - | - | 268 | 268 |
| | AF-B3DFQ1-F1 | A | 1.77 | 0.73 | 67% | 198 | 213 | 213 |

| | | | | | | | | |
|---|---|---|---|---|---|---|---|---|
| **PB0228** | P16070 | | | | | | | |
| | A0A8M6Z4J5 | | | | | | | |
| **PB0229** | P39060 | | | | | | | |
| | A0A8M9PS88 | | | | | | | |
| **PB0230** | No danio | | | | | | | |
| **PB0231** | AF-P23142-F1 | A | - | - | - | - | 703 | 703 |
| | AF-B3DH18-F1 | A | 3.5 | 0.54 | 64% | 371 | 681 | 681 |
| **PB0232** | AF-P02671-F1 | A | - | - | - | - | 866 | 866 |
| | AF-B8A5L6-F1 | A | 2.81 | 0.29 | 58% | 239 | 684 | 684 |
| **PB0233** | AF-P06396-F1 | A | - | - | - | - | 782 | 782 |
| | AF-F1QUM3-F1 | A | 5.55 | 0.71 | 61% | 478 | 729 | 729 |
| **PB0234** | AF-P08648-F1 | A | - | - | - | - | 1049 | 1049 |
| | AF-Q5I2A9-F1 | A | 2.84 | 0.89 | 54% | 924 | 1053 | 1053 |
| **PB0235** | AF-P06756-F1 | A | - | - | - | - | 1048 | 1048 |
| | AF-Q3LTM3-F1 | A | 3.34 | 0.86 | 59% | 857 | 1045 | 1045 |
| **PB0236** | P05556 | | | | | | | |
| | A0A8M6YW86 | | | | | | | |
| **PB0237** | AF-P13796-F1 | A | - | - | - | - | 627 | 627 |
| | AF-A0A140LGK1-F1 | A | 2.3 | 0.9 | 80% | 549 | 624 | 624 |
| **PB0238** | AF-P00747-F1 | A | - | - | - | - | 810 | 810 |
| | AF-F1Q890-F1 | A | 5.46 | 0.76 | 50% | 483 | 818 | 818 |
| **PB0239** | AF-Q13162-F1 | A | - | - | - | - | 271 | 271 |
| | AF-A3KP44-F1 | A | 1.51 | 0.77 | 89% | 211 | 260 | 260 |
| **PB0240** | P36952 | | | | | | | |
| | AAI62112.1 | | | | | | | |
| **PB0241** | AF-A1X283-F1 | A | - | - | - | - | 911 | 911 |
| | AF-A0A0R4IMX7-F 1 | A | 7.89 | 0.33 | 35% | 171 | 880 | 880 |
| **PB0242** | AF-P02766-F1 | A | - | - | - | - | 147 | 147 |
| | AF-B8JLL8-F1 | A | 2.72 | 0.83 | 47% | 121 | 149 | 149 |
| **PB0243** | P13611 | | | | | | | |
| **PB0244** | Q15942 | | | | | | | |
| | A0A8M9NZB7 | | | | | | | |
| **PB0245** | AF-P35222-F1 | A | - | - | - | - | 781 | 781 |
| | AF-A0A5H1ZRJ2-F1 | A | 3.22 | 0.79 | 94% | 584 | 789 | 789 |

| | | | | | | | | |
|---|---|---|---|---|---|---|---|---|
| **PB0246** | AF-P63000-F1 | A | - | - | - | - | 192 | 192 |
| | AF-Q7ZSZ9-F1 | A | 0.38 | 1 | 98% | 192 | 192 | 192 |
| **PB0247** | AF-Q96AC1-F1 | A | - | - | - | - | 680 | 680 |
| | AF-A0A2R8QN11-F1 | A | 3.92 | 0.71 | 80% | 455 | 695 | 695 |
| **PB0248** | P18206 | | | | | | | |
| | A0A8M2B3M1 | | | | | | | |
| **PB0249** | No Danio | | | | | | | |
| **PB0250** | No Danio | | | | | | | |
| **PB0251** | AF-P12644-F1 | A | - | - | - | - | 408 | 408 |
| | AF-O57574-F1 | A | 2.66 | 0.84 | 67% | 337 | 400 | 400 |
| **PB0252** | AF-Q2MKA7-F1 | A | - | - | - | - | 263 | 263 |
| | AF-Q6DHR0-F1 | A | 5.08 | 0.48 | 43% | 115 | 261 | 261 |
| **PB0253** | AF-Q6UXX9-F1 | A | - | - | - | - | 243 | 243 |
| | AF-F1Q5C9-F1 | A | 3.21 | 0.54 | 60% | 130 | 248 | 248 |
| **PB0254** | AF-Q9BXY4-F1 | A | - | - | - | - | 272 | 272 |
| | AF-Q5R328-F1 | A | 3.3 | 0.51 | 48% | 139 | 317 | 317 |
| **PB0255** | AF-Q2I0M5-F1 | A | - | - | - | - | 234 | 234 |
| | AF-F1Q5C9-F1 | A | 3.47 | 0.51 | 45% | 123 | 248 | 248 |

**Table S4.**
Pairwise sequences and structure conservation between *D. rerio and H. sapiens* homologs Pairwise structure alignment.

| | | Protein structure conservation * | | |
|---|---|---|---|---|
| | | **Low** | **Medium** | **High** |
| **Amino acids sequence conservation** ** | **Low** | BSP<br>Osteopontin | Fetuin A<br>Fetuin B ***<br>MATN3 | |
| | **Medium** | | RSPO1 *** | |
| | **High** | | | Osteonectin<br>Annexin A1<br>ALP<br>TGFβ-3 |

* *Base on the pairwise structure alignment*
** *Based on the percentage of identity*
****Fetuin B: Medium (RMSD), High (TM-Score); RSPO1: Medium (RMSD), Low (TM-Score)*

**Table S5.**
Ratios of non-synonymous (dN) and synonymous (dS) substitutions at different levels of the species tree (Fig. 1c) that includes *Danio rerio*, *Xenopus laevis*, *Alligator sinensis*, *Mus musculus*, and *Homo sapiens*.

| Species tree | | PB0001 | PB0006 | PB0059 | PB0085 | PB0092 | PB0102 | PB0195 | PB0201 | PB0249 | PB0252 |
|---|---|---|---|---|---|---|---|---|---|---|---|
| Branch A | Branch B | Fetuin A | Osteonectin | Annexin A1 | Fetuin B | ALP | Osteopontin | MATN3 | TGFβ-3 | BSP | RSPO1 |
| *M. musculus* | *H. sapiens* | 0.675 | 0.095 | 0.13 | 0.535 | 0.09 | 0.395 | 0.185 | 0.03 | 0.305 | 0.155 |
| *A. sinensis* | *M. musculus*<br>*H. sapiens* | 0.63 | 0.16 | 0.16 | **1.23** | 0.17 | 0.74 | 0.43 | 0.20 | 0.43 | 0.27 |
| *X. laevis* | *M. musculus*<br>*H. sapiens*<br>*A. sinensis* | 0.18 | 0.08 | 0.18 | 0.97 | 0.14 | Not in *X. laevis* | **2.66** | 0.37 | Not in *X. laevis* | 0.13 |
| *D. rerio* | *M.musculus*<br>*H. sapiens*<br>*A. sinensis*<br>*X. laevis* | **1.10** | 0.14 | 0.39 | **1.17** | 0.24 | **1.37** | 0.38 | 0.28 | Not in *D. rerio* | 0.27 |

*[dN/dS > 1] = positive selection (in red)*

**Table S6.**
Ratios of non-synonymous (dN) and synonymous (dS) substitutions between *Pan troglodytes* vs *Homo sapiens*.

| Phylobone Code | Protein | dN/dS |
|---|---|---|
| **PB0001** | **Fetuin A** | **2.37** |
| **PB0006** | Osteonectin | 0.00 |
| **PB0059** | Annexin A1 | 0.05 |
| **PB0085** | Fetuin B | 0.46 |
| **PB0092** | ALP | 0.05 |
| **PB0102** | **Osteopontin** | **2.07** |
| **PB0195** | MATN3 | 0.35 |
| **PB0201** | TGF-B3 | 0.00 |
| **PB0249** | BSP | 0.25 |
| **PB0252** | RSPO1 | 0.84 |

*[dN/dS > 1] = positive selection (in red)*

**Table S7.**

Gene copy numbers of selected proteins.

| Phylobone Code | Protein | Group | Gene | *Danio rerio* (GCF_000002035.6) | Homo sapiens (GCF_000001405.40) |
|---|---|---|---|---|---|
| PB0001 | Fetuin A | Group i | AHSG | 2 | 1 |
| PB0085 | Fetuin B | Group i | FETUB | 1 | 2 |
| PB0102 | Osteopontin | Group i | SPP1 | 3 | 3 |
| PB0195 | MATN3 | Group i | MATN3 | 2 | 1 |
| PB0249 | BSP | Group i | IBSP | 0 | 1 |
| PB0006 | Osteonectin | Group iii | SPARC | 6 | 7 |
| PB0092 | ALP | Group iii | ALPL | 1 | 1 |
| PB0201 | TGFB3 | Group iii | TGFB3 | 1 | 1 |
| PB0059 | Annexin A1 | Grup iv | ANXA1 | 7 | 5 |
| PB0252 | RSPO1 | Grup iv | RSPO1 | 1 | 1 |

*Group i: positive selections and mechanical stimulation, group ii: positive selection and no mechanical stimulation, group iii: purifying selection and mechanical stimulation, and group iv: purifying selection and no mechanical stimulation. Gene copy numbers obtained from NCBI Comparative Genome Viewer (CGV) (2).*

**Table S8.**
Codons usage biases in candidate proteins.

| Phylobone Code | Protein | CAI | eCAI (0.95) | Signif. vs. eCAI | Human genome CAI (Mean ± SD | Signif. vs. Human genome |
|---|---|---|---|---|---|---|
| **PB0006** | **Osteonectin** | **0.899** | 0.815 | * | 0.430 ± 0.130 | * |
| **PB0059** | **Annexin A1** | **0.769** | 0.726 | * | | * |
| **PB0092** | **ALP** | **0.843** | 0.786 | * | | * |
| **PB0201** | **TGFB3** | **0.855** | 0.788 | * | | * |
| **PB0102** | **Osteopontin** | **0.817** | 0.770 | * | | * |
| **PB0252** | **RSPO1** | **0.864** | 0.795 | * | | * |
| **PB0001** | **Fetuin A** | **0.809** | 0.755 | * | | * |
| **PB0085** | **Fetuin B** | **0.794** | 0.747 | * | | * |
| **PB0195** | **MATN3** | **0.801** | 0.765 | * | | * |
| **PB0249** | **BSP** | **0.801** | 0.802 | n.s. | | * |

*Codons adaptation index calculated with CAIcal (3,4) using the codons usage references table of ribosomal protein genes in human, obtained from (5).*

**Table S9.**
Gene expression by anatomical locations.

| Phylobone Code | Protein | Anatomical entity 1 | | Anatomical entity 2 | | Anatomical entity 3 | | Source |
|---|---|---|---|---|---|---|---|---|
| **PB0006** | **Osteonectin** | UBERON:0000979 tibia | 99.98 | CL:0002255 stromal cell of endometrium | 99.97 | UBERON:000826 periodontal ligament | 99.95 | https://www.bgee.org/gene/ENSG00000113140 |
| **PB0059** | **Annexin A1** | UBERON:0000167 oral cavity | 99.96 | UBERON:0000355 pharyngeal mucosa | 99.96 | UBERON:0035834 lower esophagus mucosa | 99.96 | https://www.bgee.org/gene/ENSG00000135046 |
| **PB0092** | **ALP** | UBERON:0001233 right adrenal gland | 96.30 | UBERON:0035827 right adrenal gland cortex | 96.61 | UBERON:0035825 left adrenal gland cortex | 95.41 | https://www.bgee.org/gene/ENSG00000162551 |
| **PB0201** | **TGFB3** | UBERON:0007318 saphenous vein | 96.03 | UBERON:0000458 endocervix | 95.69 | UBERON:0002110 gall bladder | 94.28 | https://www.bgee.org/gene/ENSG00000119699 |
| **PB0102** | **Osteopontin** | UBERON:0002110 gall bladder | 99.93 | UBERON:0002702 middle frontal gyrus | 99.93 | UBERON:0005363 inferior vagus X ganglion | 99.87 | https://www.bgee.org/gene/ENSG00000118785 |
| **PB0252** | **RSPO1** | UBERON:0000458 endocervix | 89.1 | UBERON:0009853 body of uterus | 88.97 | UBERON:0001304 germinal epithelium of ovary | 88.85 | https://www.bgee.org/gene/ENSG00000169218 |
| **PB0001** | **Fetuin A** | UBERON:0002107 liver | 99.83 | UBERON:0001114 right lobe of liver | 99.78 | UBERON:0000473 male germ line stem cell (sensu Vertebrata) *in* testis | 88.14 | https://www.bgee.org/gene/ENSG00000145192 |
| **PB0085** | **Fetuin B** | UBERON:0001114 right lobe of liver | 95.17 | UBERON:0000473 male germ line stem cell (sensu Vertebrata) *in* testis | 94.12 | UBERON:0035834 lower esophagus mucosa | 79.06 | https://www.bgee.org/gene/ENSG00000090512 |
| **PB0195** | **MATN3** | UBERON:0000979 tibia | 99.12 | UBERON:0002418 cartilage tissue | 89.97 | UBERON:0000991 primordial germ cell *in* gonad | 86.17 | https://www.bgee.org/gene/ENSG00000132031 |
| **PB0249** | **BSP** | UBERON:0000979 tibia | 99.98 | UBERON:0008266 periodontal ligament | 97.16 | UBERON:0002483 trabecular bone tissue | 95.74 | https://www.bgee.org/gene/ENSG00000029559 |

Top 3 anatomical entities. For more results click on the source link.

**Table S10.**
Protein interactions

| Phylobone Code | Protein | |
|---|---|---|
| **PB0006** | **Osteonectin (SPARC)** | 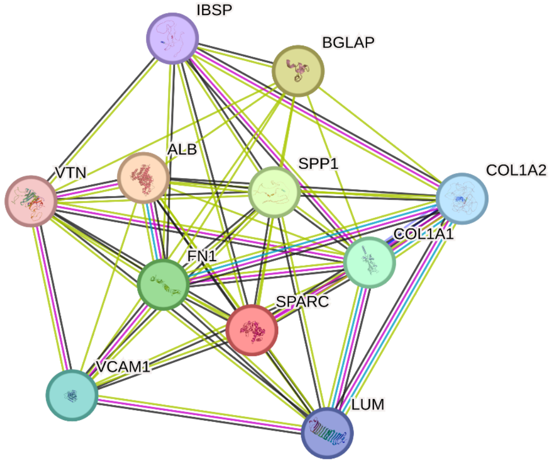 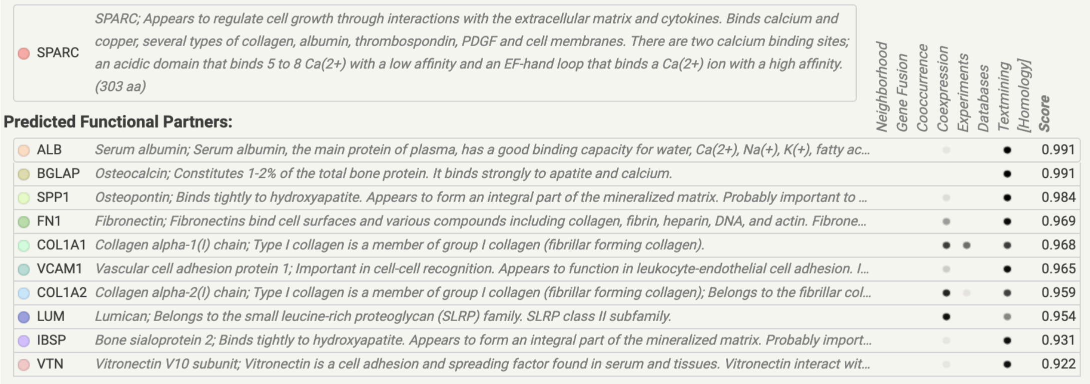 [https://string-db.org/network/9606.ENSP00000231061](https://string-db.org/network/9606.ENSP00000231061) |

SPARC — SPARC; Appears to regulate cell growth through interactions with the extracellular matrix and cytokines. Binds calcium and copper, several types of collagen, albumin, thrombospondin, PDGF and cell membranes. There are two calcium binding sites; an acidic domain that binds 5 to 8 Ca(2+) with a low affinity and an EF-hand loop that binds a Ca(2+) ion with a high affinity. (303 aa)

**Predicted Functional Partners:**

| | | Neighborhood | Gene Fusion | Cooccurrence | Coexpression | Experiments | Databases | Textmining | [Homology] | Score |
|---|---|---|---|---|---|---|---|---|---|---|
| ALB | Serum albumin; Serum albumin, the main protein of plasma, has a good binding capacity for water, Ca(2+), Na(+), K(+), fatty ac... | | | | • | | | • | | 0.991 |
| BGLAP | Osteocalcin; Constitutes 1-2% of the total bone protein. It binds strongly to apatite and calcium. | | | | | | | • | | 0.991 |
| SPP1 | Osteopontin; Binds tightly to hydroxyapatite. Appears to form an integral part of the mineralized matrix. Probably important to ... | | | | • | | | • | | 0.984 |
| FN1 | Fibronectin; Fibronectins bind cell surfaces and various compounds including collagen, fibrin, heparin, DNA, and actin. Fibrone... | | | | • | | | • | | 0.969 |
| COL1A1 | Collagen alpha-1(I) chain; Type I collagen is a member of group I collagen (fibrillar forming collagen). | | | | • | • | | • | | 0.968 |
| VCAM1 | Vascular cell adhesion protein 1; Important in cell-cell recognition. Appears to function in leukocyte-endothelial cell adhesion. I... | | | | • | | | • | | 0.965 |
| COL1A2 | Collagen alpha-2(I) chain; Type I collagen is a member of group I collagen (fibrillar forming collagen); Belongs to the fibrillar col... | | | | • | • | | • | | 0.959 |
| LUM | Lumican; Belongs to the small leucine-rich proteoglycan (SLRP) family. SLRP class II subfamily. | | | | • | | | • | | 0.954 |
| IBSP | Bone sialoprotein 2; Binds tightly to hydroxyapatite. Appears to form an integral part of the mineralized matrix. Probably import... | | | | • | | | • | | 0.931 |
| VTN | Vitronectin V10 subunit; Vitronectin is a cell adhesion and spreading factor found in serum and tissues. Vitronectin interact wit... | | | | • | | | • | | 0.922 |

<table>
<tr>
<td>PB0059</td>
<td>Annexin A1</td>
<td>
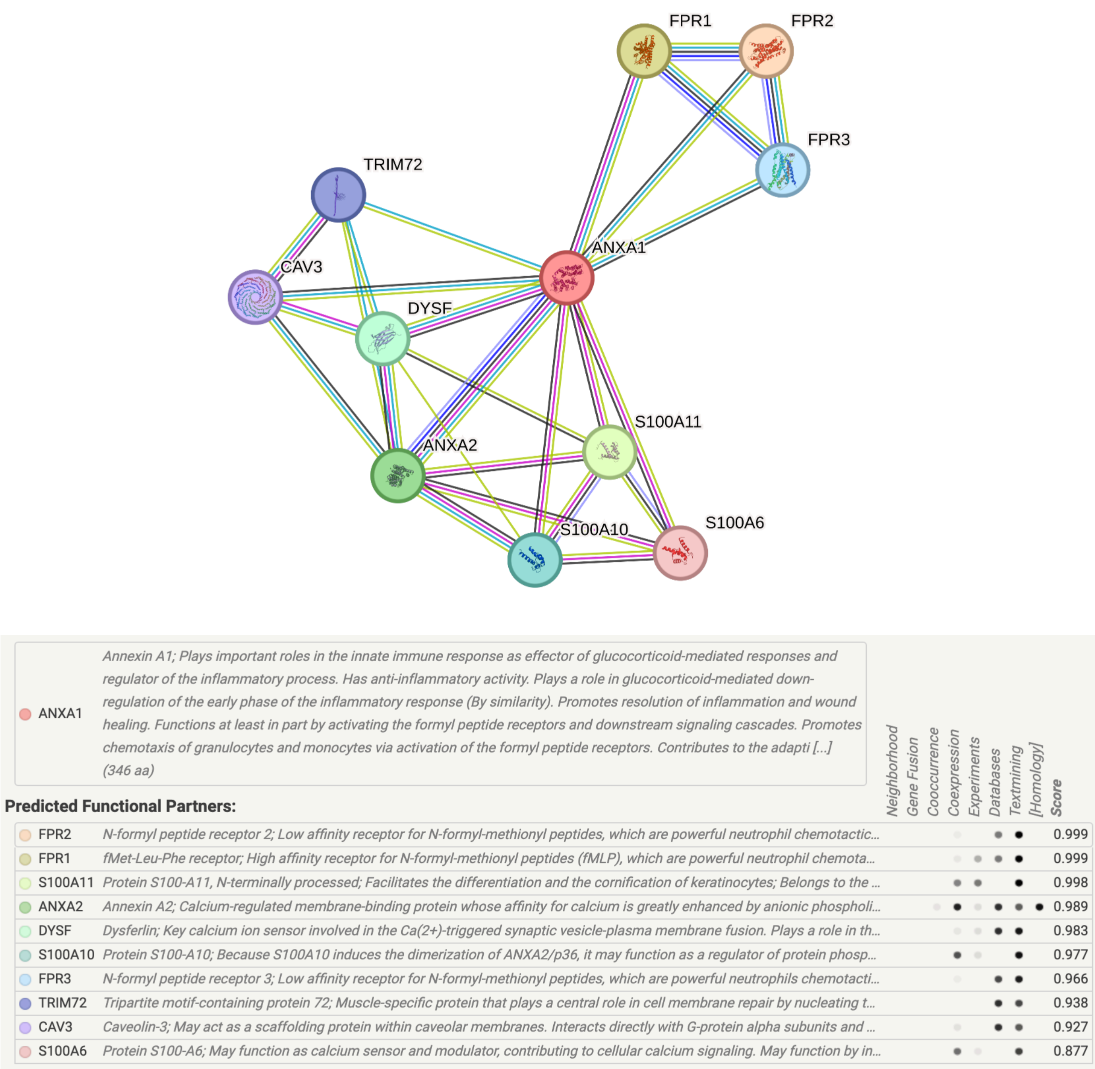

<br>
https://string-db.org/network/9606.ENSP00000366109
</td>
</tr>
</table>

| | | |
|---|---|---|
| PB0092 | ALP | see below |

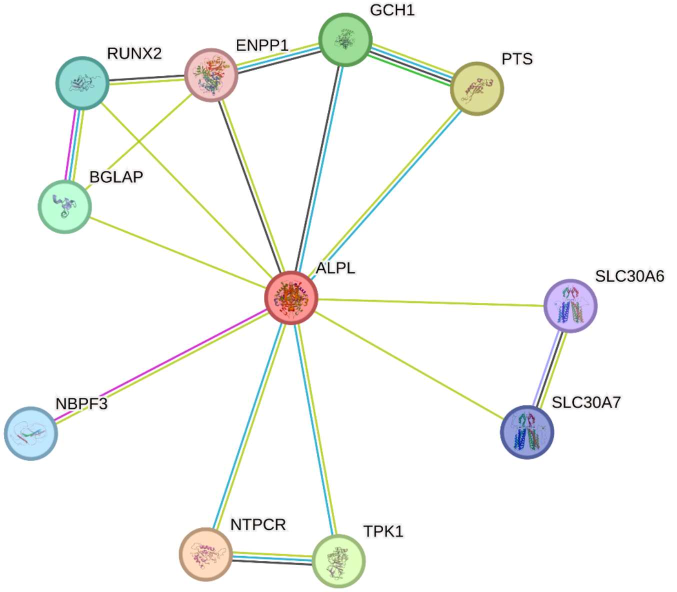

| | |
|---|---|
| ALPL | *Alkaline phosphatase, tissue-nonspecific isozyme; This isozyme plays a key role in skeletal mineralization by regulating levels of diphosphate (PPi); Belongs to the alkaline phosphatase family. (524 aa)* |

**Predicted Functional Partners:**

| | | Score |
|---|---|---|
| NTPCR | *Cancer-related nucleoside-triphosphatase; Has nucleotide phosphatase activity towards ATP, GTP, CTP, TTP and UTP. Hydroly...* | 0.905 |
| PTS | *6-pyruvoyl tetrahydrobiopterin synthase; Involved in the biosynthesis of tetrahydrobiopterin, an essential cofactor of aromatic ...* | 0.902 |
| TPK1 | *Thiamin pyrophosphokinase 1; Catalyzes the phosphorylation of thiamine to thiamine pyrophosphate. Can also catalyze the p...* | 0.901 |
| GCH1 | *GTP cyclohydrolase 1; Positively regulates nitric oxide synthesis in umbilical vein endothelial cells (HUVECs). May be involved...* | 0.901 |
| BGLAP | *Osteocalcin; Constitutes 1-2% of the total bone protein. It binds strongly to apatite and calcium.* | 0.900 |
| RUNX2 | *Runt-related transcription factor 2; Transcription factor involved in osteoblastic differentiation and skeletal morphogenesis. E...* | 0.891 |
| NBPF3 | *Neuroblastoma breakpoint family member 3; NBPF member 3; Belongs to the NBPF family.* | 0.886 |
| SLC30A7 | *Zinc transporter 7; Seems to facilitate zinc transport from the cytoplasm into the Golgi apparatus. Partly regulates cellular zin...* | 0.886 |
| SLC30A6 | *Zinc transporter 6; Zinc-efflux transporter which allocates the cytoplasmic zinc to the trans-Golgi network (TGN) as well as th...* | 0.885 |
| ENPP1 | *Ectonucleotide pyrophosphatase/phosphodiesterase family member 1, secreted form; Nucleotide pyrophosphatase that gene...* | 0.870 |

Evidence columns: Neighborhood, Gene Fusion, Cooccurrence, Coexpression, Experiments, Databases, Textmining, [Homology], Score

https://string-db.org/network/9606.ENSP00000363973

<table>
<tr>
<td>PB0201</td>
<td>TGFB3</td>
<td>
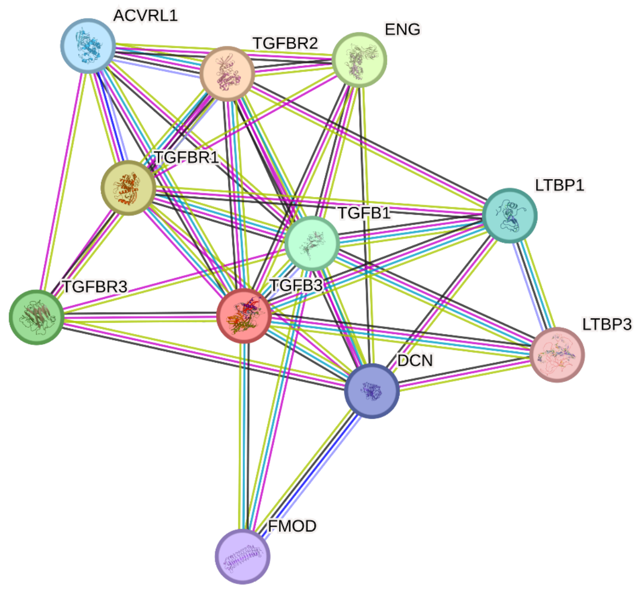

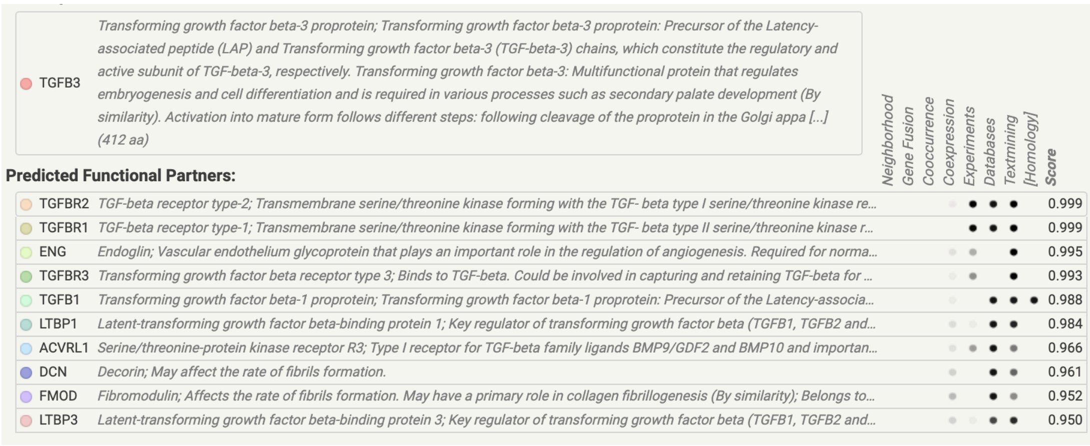
<table>
<tr><td>TGFB3</td><td>Transforming growth factor beta-3 proprotein; Transforming growth factor beta-3 proprotein: Precursor of the Latency-associated peptide (LAP) and Transforming growth factor beta-3 (TGF-beta-3) chains, which constitute the regulatory and active subunit of TGF-beta-3, respectively. Transforming growth factor beta-3: Multifunctional protein that regulates embryogenesis and cell differentiation and is required in various processes such as secondary palate development (By similarity). Activation into mature form follows different steps: following cleavage of the proprotein in the Golgi appa [...] (412 aa)</td></tr>
</table>
Predicted Functional Partners:
<table>
<tr><th></th><th></th><th>Score</th></tr>
<tr><td>TGFBR2</td><td>TGF-beta receptor type-2; Transmembrane serine/threonine kinase forming with the TGF- beta type I serine/threonine kinase re...</td><td>0.999</td></tr>
<tr><td>TGFBR1</td><td>TGF-beta receptor type-1; Transmembrane serine/threonine kinase forming with the TGF- beta type II serine/threonine kinase r...</td><td>0.999</td></tr>
<tr><td>ENG</td><td>Endoglin; Vascular endothelium glycoprotein that plays an important role in the regulation of angiogenesis. Required for norma...</td><td>0.995</td></tr>
<tr><td>TGFBR3</td><td>Transforming growth factor beta receptor type 3; Binds to TGF-beta. Could be involved in capturing and retaining TGF-beta for ...</td><td>0.993</td></tr>
<tr><td>TGFB1</td><td>Transforming growth factor beta-1 proprotein; Transforming growth factor beta-1 proprotein: Precursor of the Latency-associa...</td><td>0.988</td></tr>
<tr><td>LTBP1</td><td>Latent-transforming growth factor beta-binding protein 1; Key regulator of transforming growth factor beta (TGFB1, TGFB2 and...</td><td>0.984</td></tr>
<tr><td>ACVRL1</td><td>Serine/threonine-protein kinase receptor R3; Type I receptor for TGF-beta family ligands BMP9/GDF2 and BMP10 and importan...</td><td>0.966</td></tr>
<tr><td>DCN</td><td>Decorin; May affect the rate of fibrils formation.</td><td>0.961</td></tr>
<tr><td>FMOD</td><td>Fibromodulin; Affects the rate of fibrils formation. May have a primary role in collagen fibrillogenesis (By similarity); Belongs to...</td><td>0.952</td></tr>
<tr><td>LTBP3</td><td>Latent-transforming growth factor beta-binding protein 3; Key regulator of transforming growth factor beta (TGFB1, TGFB2 and...</td><td>0.950</td></tr>
</table>
(Evidence columns: Neighborhood, Gene Fusion, Cooccurrence, Coexpression, Experiments, Databases, Textmining, [Homology])
<br>
https://string-db.org/network/9606.ENSP00000238682
</td>
</tr>
<tr>
<td>PB0102</td>
<td>Osteopontin</td>
<td>
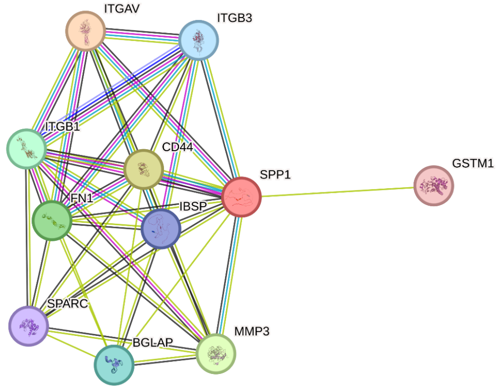

</td>
</tr>
</table>

| | |
|---|---|
| SPP1 | *Osteopontin; Binds tightly to hydroxyapatite. Appears to form an integral part of the mineralized matrix. Probably important to cell-matrix interaction. (314 aa)* |

**Predicted Functional Partners:**

| Gene | Description | Score |
|---|---|---|
| ITGAV | *Integrin alpha-V heavy chain; The alpha-V (ITGAV) integrins are receptors for vitronectin, cytotactin, fibronectin, fibrinogen, lami...* | 0.999 |
| CD44 | *CD44 antigen; Cell-surface receptor that plays a role in cell-cell interactions, cell adhesion and migration, helping them to sense ...* | 0.999 |
| MMP3 | *Stromelysin-1; Can degrade fibronectin, laminin, gelatins of type I, III, IV, and V; collagens III, IV, X, and IX, and cartilage proteogly...* | 0.997 |
| FN1 | *Fibronectin; Fibronectins bind cell surfaces and various compounds including collagen, fibrin, heparin, DNA, and actin. Fibronec...* | 0.996 |
| ITGB1 | *Integrin beta-1; Integrins alpha-1/beta-1, alpha-2/beta-1, alpha-10/beta-1 and alpha-11/beta-1 are receptors for collagen. Integri...* | 0.996 |
| BGLAP | *Osteocalcin; Constitutes 1-2% of the total bone protein. It binds strongly to apatite and calcium.* | 0.994 |
| ITGB3 | *Integrin beta-3; Integrin alpha-V/beta-3 (ITGAV:ITGB3) is a receptor for cytotactin, fibronectin, laminin, matrix metalloproteinase...* | 0.994 |
| IBSP | *Bone sialoprotein 2; Binds tightly to hydroxyapatite. Appears to form an integral part of the mineralized matrix. Probably import...* | 0.991 |
| SPARC | *SPARC; Appears to regulate cell growth through interactions with the extracellular matrix and cytokines. Binds calcium and cop...* | 0.984 |
| GSTM1 | *Glutathione S-transferase Mu 1; Conjugation of reduced glutathione to a wide number of exogenous and endogenous hydropho...* | 0.981 |

https://string-db.org/network/9606.ENSP00000378517

**PB0252** | **RSPO1**

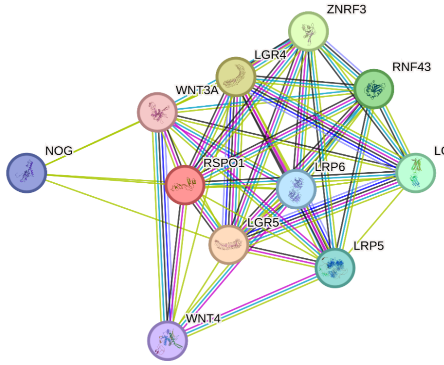

| | |
|---|---|
| RSPO1 | *R-spondin-1; Activator of the canonical Wnt signaling pathway by acting as a ligand for LGR4-6 receptors. Upon binding to LGR4-6 (LGR4, LGR5 or LGR6), LGR4-6 associate with phosphorylated LRP6 and frizzled receptors that are activated by extracellular Wnt receptors, triggering the canonical Wnt signaling pathway to increase expression of target genes. Also regulates the canonical Wnt/beta-catenin- dependent pathway and non-canonical Wnt signaling by acting as an inhibitor of ZNRF3, an important regulator of the Wnt signaling pathway. Acts as a ligand for frizzled FZD8 and LRP6. May neg [...] (263 aa)* |

**Predicted Functional Partners:**

| Gene | Description | Score |
|---|---|---|
| LGR5 | *Leucine-rich repeat-containing G-protein coupled receptor 5; Receptor for R-spondins that potentiates the canonical Wnt signali...* | 0.999 |
| LGR4 | *Leucine-rich repeat-containing G-protein coupled receptor 4; Receptor for R-spondins that potentiates the canonical Wnt signali...* | 0.999 |
| ZNRF3 | *E3 ubiquitin-protein ligase ZNRF3; E3 ubiquitin-protein ligase that acts as a negative regulator of the Wnt signaling pathway by ...* | 0.999 |
| RNF43 | *E3 ubiquitin-protein ligase RNF43; E3 ubiquitin-protein ligase that acts as a negative regulator of the Wnt signaling pathway by ...* | 0.999 |
| LGR6 | *Leucine-rich repeat-containing G-protein coupled receptor 6; Receptor for R-spondins that potentiates the canonical Wnt signali...* | 0.998 |
| LRP5 | *Low-density lipoprotein receptor-related protein 5; Acts as a coreceptor with members of the frizzled family of seven-transmem...* | 0.961 |
| LRP6 | *Low-density lipoprotein receptor-related protein 6; Component of the Wnt-Fzd-LRP5-LRP6 complex that triggers beta-catenin si...* | 0.959 |
| NOG | *Noggin; Inhibitor of bone morphogenetic proteins (BMP) signaling which is required for growth and patterning of the neural tub...* | 0.925 |
| WNT4 | *Protein Wnt-4; Ligand for members of the frizzled family of seven transmembrane receptors (Probable). Plays an important rol...* | 0.923 |
| WNT3A | *Protein Wnt-3a; Ligand for members of the frizzled family of seven transmembrane receptors (Probable). Functions in the cano...* | 0.903 |

https://string-db.org/network/9606.ENSP00000348944

<table>
<tr>
<td>PB0001</td>
<td>Fetuin A</td>
<td>
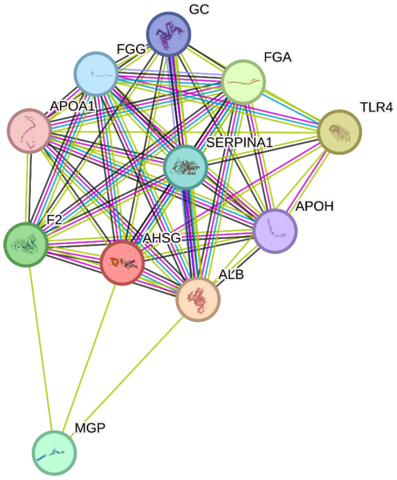

<table>
<tr><td>AHSG</td><td colspan="10">Alpha-2-HS-glycoprotein chain A; Promotes endocytosis, possesses opsonic properties and influences the mineral phase of bone. Shows affinity for calcium and barium ions; Belongs to the fetuin family. (368 aa)</td></tr>
<tr><th colspan="2">Predicted Functional Partners:</th><th>Neighborhood</th><th>Gene Fusion</th><th>Cooccurrence</th><th>Coexpression</th><th>Experiments</th><th>Databases</th><th>Textmining</th><th>[Homology]</th><th>Score</th></tr>
<tr><td>ALB</td><td>Serum albumin; Serum albumin, the main protein of plasma, has a good binding capacity for water, Ca(2+), Na(+), K(+), fatty ...</td><td></td><td></td><td></td><td>●</td><td>●</td><td>●</td><td>●</td><td></td><td>0.999</td></tr>
<tr><td>TLR4</td><td>Toll-like receptor 4; Cooperates with LY96 and CD14 to mediate the innate immune response to bacterial lipopolysaccharide ...</td><td></td><td></td><td></td><td></td><td>●</td><td></td><td>●</td><td></td><td>0.988</td></tr>
<tr><td>FGA</td><td>Fibrinogen alpha chain; Cleaved by the protease thrombin to yield monomers which, together with fibrinogen beta (FGB) and ...</td><td></td><td></td><td></td><td>●</td><td>●</td><td>●</td><td>●</td><td></td><td>0.984</td></tr>
<tr><td>F2</td><td>Activation peptide fragment 1; Thrombin, which cleaves bonds after Arg and Lys, converts fibrinogen to fibrin and activates f...</td><td></td><td></td><td></td><td>●</td><td>●</td><td></td><td>●</td><td></td><td>0.980</td></tr>
<tr><td>MGP</td><td>Matrix Gla protein; Associates with the organic matrix of bone and cartilage. Thought to act as an inhibitor of bone formation.</td><td></td><td></td><td></td><td></td><td></td><td></td><td>●</td><td></td><td>0.977</td></tr>
<tr><td>SERPINA1</td><td>Short peptide from AAT; Inhibitor of serine proteases. Its primary target is elastase, but it also has a moderate affinity for pla...</td><td></td><td></td><td></td><td>●</td><td>●</td><td>●</td><td>●</td><td></td><td>0.971</td></tr>
<tr><td>FGG</td><td>Fibrinogen gamma chain; Together with fibrinogen alpha (FGA) and fibrinogen beta (FGB), polymerizes to form an insoluble ...</td><td></td><td></td><td></td><td>●</td><td>●</td><td>●</td><td>●</td><td></td><td>0.970</td></tr>
<tr><td>GC</td><td>Vitamin D-binding protein; Involved in vitamin D transport and storage, scavenging of extracellular G-actin, enhancement of t...</td><td></td><td></td><td></td><td>●</td><td></td><td></td><td>●</td><td></td><td>0.959</td></tr>
<tr><td>APOH</td><td>Beta-2-glycoprotein 1; Binds to various kinds of negatively charged substances such as heparin, phospholipids, and dextran ...</td><td></td><td></td><td></td><td>●</td><td></td><td></td><td>●</td><td></td><td>0.957</td></tr>
<tr><td>APOA1</td><td>Truncated apolipoprotein A-I; Participates in the reverse transport of cholesterol from tissues to the liver for excretion by pro...</td><td></td><td></td><td></td><td>●</td><td></td><td></td><td>●</td><td></td><td>0.944</td></tr>
</table>
https://string-db.org/network/9606.ENSP00000273784
</td>
</tr>
<tr>
<td>PB0085</td>
<td>Fetuin B</td>
<td>
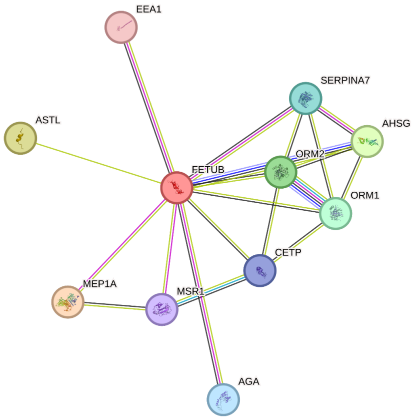

</td>
</tr>
</table>

<table>
<tr><td></td><td></td><td>

FETUB — Fetuin-B; Protease inhibitor required for egg fertilization. Required to prevent premature zona pellucida hardening before fertilization, probably by inhibiting the protease activity of ASTL, a protease that mediates the cleavage of ZP2 and triggers zona pellucida hardening (By similarity). (382 aa)

**Predicted Functional Partners:**

Neighborhood, Gene Fusion, Cooccurrence, Coexpression, Experiments, Databases, Textmining, [Homology], Score

| Partner | Description | Score |
|---|---|---|
| MEP1A | Meprin A subunit alpha. | 0.900 |
| ASTL | Astacin-like metalloendopeptidase; Oocyte-specific oolemmal receptor involved in sperm and egg adhesion and fertilization.... | 0.701 |
| AHSG | Alpha-2-HS-glycoprotein chain A; Promotes endocytosis, possesses opsonic properties and influences the mineral phase of ... | 0.659 |
| ORM2 | Alpha-1-acid glycoprotein 2; Functions as transport protein in the blood stream. Binds various hydrophobic ligands in the int... | 0.617 |
| ORM1 | Alpha-1-acid glycoprotein 1; Functions as transport protein in the blood stream. Binds various ligands in the interior of its bet... | 0.601 |
| SERPINA7 | Thyroxine-binding globulin; Major thyroid hormone transport protein in serum; Belongs to the serpin family. | 0.583 |
| AGA | N(4)-(beta-N-acetylglucosaminyl)-L-asparaginase; Cleaves the GlcNAc-Asn bond which joins oligosaccharides to the peptide... | 0.570 |
| CETP | Cholesteryl ester transfer protein; Involved in the transfer of neutral lipids, including cholesteryl ester and triglyceride, among... | 0.566 |
| MSR1 | Macrophage scavenger receptor types I and II; Membrane glycoproteins implicated in the pathologic deposition of cholester... | 0.559 |
| EEA1 | Early endosome antigen 1; Binds phospholipid vesicles containing phosphatidylinositol 3-phosphate and participates in end... | 0.556 |

https://string-db.org/network/9606.ENSP00000265029

</td></tr>
<tr><td>**PB0195**</td><td>**MATN3**</td><td>

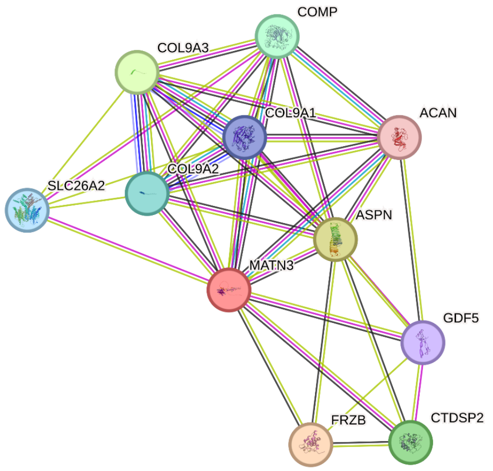

MATN3 — Matrilin-3; Major component of the extracellular matrix of cartilage and may play a role in the formation of extracellular filamentous networks. (486 aa)

**Predicted Functional Partners:**

Neighborhood, Gene Fusion, Cooccurrence, Coexpression, Experiments, Databases, Textmining, [Homology], Score

| Partner | Description | Score |
|---|---|---|
| FRZB | Secreted frizzled-related protein 3; Soluble frizzled-related proteins (sFRPS) function as modulators of Wnt signaling through ... | 0.953 |
| ASPN | Asporin; Negatively regulates periodontal ligament (PDL) differentiation and mineralization to ensure that the PDL is not ossifi... | 0.940 |
| COL9A3 | Collagen alpha-3(IX) chain; Structural component of hyaline cartilage and vitreous of the eye. | 0.929 |
| CTDSP2 | Carboxy-terminal domain RNA polymerase II polypeptide A small phosphatase 2; Preferentially catalyzes the dephosphorylati... | 0.926 |
| COMP | Cartilage oligomeric matrix protein; May play a role in the structural integrity of cartilage via its interaction with other extracell... | 0.923 |
| COL9A2 | Collagen alpha-2(IX) chain; Structural component of hyaline cartilage and vitreous of the eye; Belongs to the fibril-associated ... | 0.921 |
| SLC26A2 | Sulfate transporter; Sulfate transporter. May play a role in endochondral bone formation. | 0.907 |
| COL9A1 | Collagen alpha-1(IX) chain; Structural component of hyaline cartilage and vitreous of the eye. | 0.891 |
| GDF5 | Growth/differentiation factor 5; Growth factor involved in bone and cartilage formation. During cartilage development regulat... | 0.765 |
| ACAN | Aggrecan core protein 2; This proteoglycan is a major component of extracellular matrix of cartilagenous tissues. A major fun... | 0.730 |

https://string-db.org/network/9606.ENSP00000383894

</td></tr>
</table>

**PB0249** | **BSP**

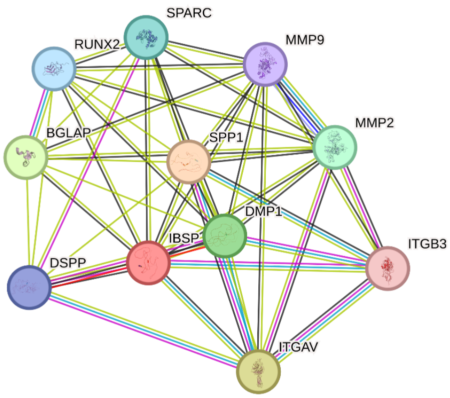

| | |
|---|---|
| IBSP | *Bone sialoprotein 2; Binds tightly to hydroxyapatite. Appears to form an integral part of the mineralized matrix. Probably important to cell-matrix interaction. Promotes Arg-Gly-Asp-dependent cell attachment. (317 aa)* |

**Predicted Functional Partners:**

| | | Neighborhood | Gene Fusion | Cooccurrence | Coexpression | Experiments | Databases | Textmining | [Homology] | Score |
|---|---|---|---|---|---|---|---|---|---|---|
| SPP1 | *Osteopontin; Binds tightly to hydroxyapatite. Appears to form an integral part of the mineralized matrix. Probably important to c...* | | | | ● | | | ● | | 0.991 |
| ITGAV | *Integrin alpha-V heavy chain; The alpha-V (ITGAV) integrins are receptors for vitronectin, cytotactin, fibronectin, fibrinogen, lamin...* | | | | ● | | ● | ● | | 0.983 |
| BGLAP | *Osteocalcin; Constitutes 1-2% of the total bone protein. It binds strongly to apatite and calcium.* | | | | ● | | | ● | | 0.971 |
| DMP1 | *Dentin matrix acidic phosphoprotein 1; May have a dual function during osteoblast differentiation. In the nucleus of undifferenti...* | | | | ● | | | ● | | 0.969 |
| MMP2 | *72 kDa type IV collagenase; Ubiquitinous metalloproteinase that is involved in diverse functions such as remodeling of the vasc...* | | | | ● | | | ● | | 0.936 |
| SPARC | *SPARC; Appears to regulate cell growth through interactions with the extracellular matrix and cytokines. Binds calcium and cop...* | | | | ● | | | ● | | 0.931 |
| RUNX2 | *Runt-related transcription factor 2; Transcription factor involved in osteoblastic differentiation and skeletal morphogenesis. Ess...* | | | | ● | | | ● | | 0.931 |
| DSPP | *Dentin sialophosphoprotein; DSP may be an important factor in dentinogenesis. DPP may bind high amount of calcium and faci...* | | | | ● | ● | | ● | | 0.930 |
| MMP9 | *67 kDa matrix metalloproteinase-9; May play an essential role in local proteolysis of the extracellular matrix and in leukocyte mi...* | | | | ● | | | ● | | 0.928 |
| ITGB3 | *Integrin beta-3; Integrin alpha-V/beta-3 (ITGAV:ITGB3) is a receptor for cytotactin, fibronectin, laminin, matrix metalloproteinase-...* | | | | | ● | ● | ● | | 0.923 |

https://string-db.org/network/9606.ENSP00000226284

**Network legend**

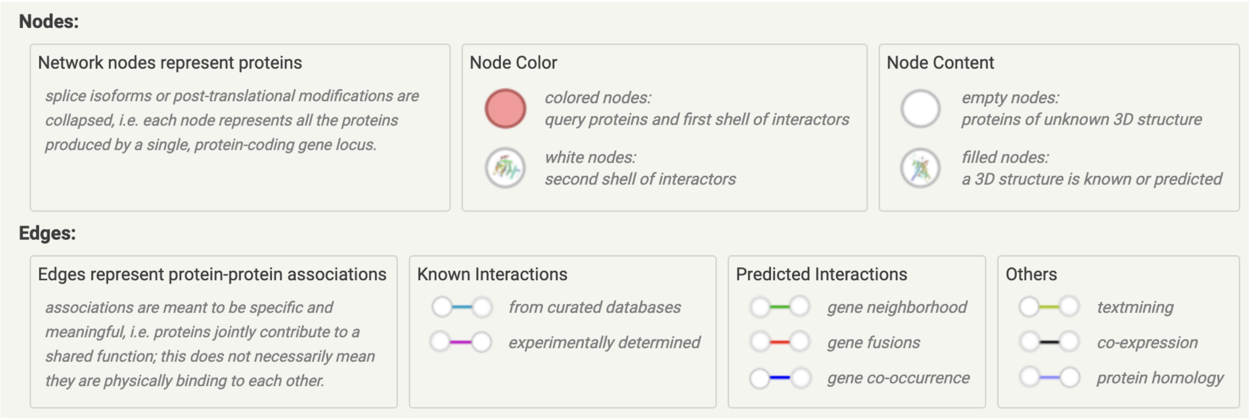

Putative protein-protein interactions are obtained from String v.12.0 at https://string-db.org (6)

**Table S11.**
Gene ontology annotations of the selected proteins.

| Protein (Name) | Length | Gene Ontology |
|---|---|---|
| PB0249 (BSP)* | 317 | GO:0003674 (enables) molecular_function<br>GO:0005178 (enables) integrin binding<br>GO:0005515 (enables) protein binding<br>GO:0001503 (involved in) ossification<br>GO:0001649 (involved in) osteoblast differentiation<br>GO:0007155 (involved in) cell adhesion<br>GO:0030198 (involved in) extracellular matrix organization<br>GO:0030282 (involved in) bone mineralization<br>GO:0031214 (involved in) biomineral tissue development<br>GO:0045785 (involved in) positive regulation of cell adhesion<br>GO:0071363 (involved in) cellular response to growth factor stimulus<br>GO:0005576 (located in) extracellular region<br>GO:0005615 (located in) extracellular space<br>GO:0016020 (located in) membrane<br>GO:0031982 (located in) vesicle |
| PB0102 (Osteopontin) | 344 | GO:0005125 (enables) cytokine activity<br>GO:0005178 (enables) integrin binding<br>GO:0005515 (enables) protein binding<br>GO:0050840 (enables) extracellular matrix binding<br>GO:0001503 (involved in) ossification<br>GO:0001649 (involved in) osteoblast differentiation<br>GO:0006710 (involved in) androgen catabolic process<br>GO:0006954 (involved in) inflammatory response<br>GO:0007155 (involved in) cell adhesion<br>GO:0007165 (involved in) signal transduction<br>GO:0007566 (involved in) embryo implantation<br>GO:0010033 (involved in) response to organic substance<br>GO:0030154 (involved in) cell differentiation<br>GO:0031214 (involved in) biomineral tissue development<br>GO:0033280 (involved in) response to vitamin D<br>GO:0045780 (involved in) positive regulation of bone resorption<br>GO:0045893 (involved in) positive regulation of transcription, DNA-templated<br>GO:0046697 (involved in) decidualization<br>GO:0048545 (involved in) response to steroid hormone<br>GO:0048685 (involved in) negative regulation of collateral sprouting of intact axon in response to injury<br>GO:0071394 (involved in) cellular response to testosterone stimulus<br>GO:2000866 (involved in) positive regulation of estradiol secretion<br>GO:0005615 (is active in) extracellular space<br>GO:0005576 (located in) extracellular region<br>GO:0005788 (located in) endoplasmic reticulum lumen<br>GO:0005794 (located in) Golgi apparatus<br>GO:0031982 (located in) vesicle<br>GO:0042995 (located in) cell projection<br>GO:0048471 (located in) perinuclear region of cytoplasm<br>GO:0070062 (located in) extracellular exosome |
| PB0085 (Fetuin B) | 507 | GO:0003674 (enables) molecular_function<br>GO:0004857 (enables) enzyme inhibitor activity<br>GO:0004866 (enables) endopeptidase inhibitor activity<br>GO:0004869 (enables) cysteine-type endopeptidase inhibitor activity |

| | | |
|---|---|---|
| | | GO:0005515 (enables) protein binding<br>GO:0008191 (enables) metalloendopeptidase inhibitor activity<br>GO:0030414 (enables) peptidase inhibitor activity<br>GO:0007338 (involved in) single fertilization<br>GO:0007339 (involved in) binding of sperm to zona pellucida<br>GO:0008150 (involved in) biological_process<br>GO:0010466 (involved in) negative regulation of peptidase activity<br>GO:0010951 (involved in) negative regulation of endopeptidase activity<br>GO:0005576 (located in) extracellular region<br>GO:0005615 (located in) extracellular space<br>GO:0070062 (located in) extracellular exosome |
| PB0001 (Fetuin A, AHSG) | 418 | GO:0004866 (enables) endopeptidase inhibitor activity<br>GO:0004869 (enables) cysteine-type endopeptidase inhibitor activity<br>GO:0019210 (enables) kinase inhibitor activity<br>GO:0001501 (involved in) skeletal system development<br>GO:0001503 (involved in) ossification<br>GO:0006907 (involved in) pinocytosis<br>GO:0006953 (involved in) acute-phase response<br>GO:0010951 (involved in) negative regulation of endopeptidase activity<br>GO:0030500 (involved in) regulation of bone mineralization<br>GO:0030502 (involved in) negative regulation of bone mineralization<br>GO:0046627 (involved in) negative regulation of insulin receptor signaling pathway<br>GO:0050727 (involved in) regulation of inflammatory response<br>GO:0050766 (involved in) positive regulation of phagocytosis<br>GO:0031012 (is active in) extracellular matrix<br>GO:0005576 (located in) extracellular region<br>GO:0005615 (located in) extracellular space<br>GO:0005788 (located in) endoplasmic reticulum lumen<br>GO:0005794 (located in) Golgi apparatus<br>GO:0031093 (located in) platelet alpha granule lumen<br>GO:0034774 (located in) secretory granule lumen<br>GO:0062023 (located in) collagen-containing extracellular matrix<br>GO:0070062 (located in) extracellular exosome<br>GO:0072562 (located in) blood microparticle |
| PB0195 (MATN3) | 489 | GO:0005201 (enables) extracellular matrix structural constituent<br>GO:0005509 (enables) calcium ion binding<br>GO:0005515 (enables) protein binding<br>GO:0001501 (involved in) skeletal system development<br>GO:0030198 (involved in) extracellular matrix organization<br>GO:0051216 (involved in) cartilage development<br>GO:0005576 (located in) extracellular region<br>GO:0005788 (located in) endoplasmic reticulum lumen<br>GO:0031012 (located in) extracellular matrix<br>GO:0062023 (located in) collagen-containing extracellular matrix<br>GO:0120216 (part of) matrilin complex |
| PB0252 (RSPO1) | 272 | GO:0001664 (enables) G protein-coupled receptor binding<br>GO:0005102 (enables) signaling receptor binding<br>GO:0005515 (enables) protein binding<br>GO:0008201 (enables) heparin binding<br>GO:0001934 (involved in) positive regulation of protein phosphorylation<br>GO:0002090 (involved in) regulation of receptor internalization<br>GO:0016055 (involved in) Wnt signaling pathway<br>GO:0030177 (involved in) positive regulation of Wnt signaling pathway<br>GO:0050896 (involved in) response to stimulus<br>GO:0090263 (involved in) positive regulation of canonical Wnt signaling pathway<br>GO:0005576 (located in) extracellular region<br>GO:0005634 (located in) nucleus |
| PB0092 (ALP) | 564 | GO:0003824 (enables) catalytic activity<br>GO:0004035 (enables) alkaline phosphatase activity |

| | | |
|---|---|---|
| | | GO:0004427 (enables) inorganic diphosphatase activity<br>GO:0005509 (enables) calcium ion binding<br>GO:0005515 (enables) protein binding<br>GO:0016462 (enables) pyrophosphatase activity<br>GO:0016787 (enables) hydrolase activity<br>GO:0016791 (enables) phosphatase activity<br>GO:0016887 (enables) ATP hydrolysis activity<br>GO:0033883 (enables) pyridoxal phosphatase activity<br>GO:0043262 (enables) adenosine-diphosphatase activity<br>GO:0046872 (enables) metal ion binding<br>GO:0050187 (enables) phosphoamidase activity<br>GO:0052732 (enables) phosphoethanolamine phosphatase activity<br>GO:0001501 (involved in) skeletal system development<br>GO:0001649 (involved in) osteoblast differentiation<br>GO:0001958 (involved in) endochondral ossification<br>GO:0003006 (involved in) developmental process involved in reproduction<br>GO:0016311 (involved in) dephosphorylation<br>GO:0030282 (involved in) bone mineralization<br>GO:0031214 (involved in) biomineral tissue development<br>GO:0032496 (involved in) response to lipopolysaccharide<br>GO:0033280 (involved in) response to vitamin D<br>GO:0042822 (involved in) pyridoxal phosphate metabolic process<br>GO:0046677 (involved in) response to antibiotic<br>GO:0051384 (involved in) response to glucocorticoid<br>GO:0071407 (involved in) cellular response to organic cyclic compound<br>GO:0071529 (involved in) cementum mineralization<br>GO:0120162 (involved in) positive regulation of cold-induced thermogenesis<br>GO:0005576 (located in) extracellular region<br>GO:0005615 (located in) extracellular space<br>GO:0005739 (located in) mitochondrion<br>GO:0005758 (located in) mitochondrial intermembrane space<br>GO:0005886 (located in) plasma membrane<br>GO:0016020 (located in) membrane<br>GO:0031012 (located in) extracellular matrix<br>GO:0031225 (located in) anchored component of membrane<br>GO:0031966 (located in) mitochondrial membrane<br>GO:0065010 (located in) extracellular membrane-bounded organelle<br>GO:0070062 (located in) extracellular exosome |
| PB0059 (Annexin A1) | 346 | GO:0005884 (colocalizes with) actin filament<br>GO:0004859 (enables) phospholipase inhibitor activity<br>GO:0005102 (enables) signaling receptor binding<br>GO:0005509 (enables) calcium ion binding<br>GO:0005515 (enables) protein binding<br>GO:0005543 (enables) phospholipid binding<br>GO:0005544 (enables) calcium-dependent phospholipid binding<br>GO:0019834 (enables) phospholipase A2 inhibitor activity<br>GO:0046872 (enables) metal ion binding<br>GO:0048306 (enables) calcium-dependent protein binding<br>GO:0098641 (enables) cadherin binding involved in cell-cell adhesion<br>GO:0001780 (involved in) neutrophil homeostasis<br>GO:0002250 (involved in) adaptive immune response<br>GO:0002376 (involved in) immune system process<br>GO:0002548 (involved in) monocyte chemotaxis<br>GO:0002685 (involved in) regulation of leukocyte migration<br>GO:0006909 (involved in) phagocytosis<br>GO:0006954 (involved in) inflammatory response<br>GO:0007165 (involved in) signal transduction<br>GO:0007166 (involved in) cell surface receptor signaling pathway<br>GO:0007187 (involved in) G protein-coupled receptor signaling pathway, coupled to cyclic nucleotide second messenger<br>GO:0008360 (involved in) regulation of cell shape |

| | | |
|---|---|---|
| | | GO:0009410 (involved in) response to xenobiotic stimulus<br>GO:0010165 (involved in) response to X-ray<br>GO:0014839 (involved in) myoblast migration involved in skeletal muscle regeneration<br>GO:0018149 (involved in) peptide cross-linking<br>GO:0030073 (involved in) insulin secretion<br>GO:0030216 (involved in) keratinocyte differentiation<br>GO:0030850 (involved in) prostate gland development<br>GO:0031018 (involved in) endocrine pancreas development<br>GO:0031340 (involved in) positive regulation of vesicle fusion<br>GO:0031394 (involved in) positive regulation of prostaglandin biosynthetic process<br>GO:0031532 (involved in) actin cytoskeleton reorganization<br>GO:0032355 (involved in) response to estradiol<br>GO:0032508 (involved in) DNA duplex unwinding<br>GO:0032652 (involved in) regulation of interleukin-1 production<br>GO:0032717 (involved in) negative regulation of interleukin-8 production<br>GO:0032743 (involved in) positive regulation of interleukin-2 production<br>GO:0033031 (involved in) positive regulation of neutrophil apoptotic process<br>GO:0035924 (involved in) cellular response to vascular endothelial growth factor stimulus<br>GO:0042063 (involved in) gliogenesis<br>GO:0042102 (involved in) positive regulation of T cell proliferation<br>GO:0042127 (involved in) regulation of cell population proliferation<br>GO:0043066 (involved in) negative regulation of apoptotic process<br>GO:0043434 (involved in) response to peptide hormone<br>GO:0045087 (involved in) innate immune response<br>GO:0045627 (involved in) positive regulation of T-helper 1 cell differentiation<br>GO:0045629 (involved in) negative regulation of T-helper 2 cell differentiation<br>GO:0045920 (involved in) negative regulation of exocytosis<br>GO:0046632 (involved in) alpha-beta T cell differentiation<br>GO:0046883 (involved in) regulation of hormone secretion<br>GO:0050482 (involved in) arachidonic acid secretion<br>GO:0050727 (involved in) regulation of inflammatory response<br>GO:0070301 (involved in) cellular response to hydrogen peroxide<br>GO:0070365 (involved in) hepatocyte differentiation<br>GO:0070459 (involved in) prolactin secretion<br>GO:0070555 (involved in) response to interleukin-1<br>GO:0071385 (involved in) cellular response to glucocorticoid stimulus<br>GO:0071621 (involved in) granulocyte chemotaxis<br>GO:0090050 (involved in) positive regulation of cell migration involved in sprouting angiogenesis<br>GO:0090303 (involved in) positive regulation of wound healing<br>GO:0097350 (involved in) neutrophil clearance<br>GO:0098609 (involved in) cell-cell adhesion<br>GO:1900087 (involved in) positive regulation of G1/S transition of mitotic cell cycle<br>GO:1900138 (involved in) negative regulation of phospholipase A2 activity<br>GO:0001533 (located in) cornified envelope<br>GO:0001891 (located in) phagocytic cup<br>GO:0005576 (located in) extracellular region<br>GO:0005615 (located in) extracellular space<br>GO:0005634 (located in) nucleus<br>GO:0005654 (located in) nucleoplasm<br>GO:0005737 (located in) cytoplasm<br>GO:0005768 (located in) endosome<br>GO:0005769 (located in) early endosome<br>GO:0005829 (located in) cytosol<br>GO:0005886 (located in) plasma membrane<br>GO:0005912 (located in) adherens junction<br>GO:0005925 (located in) focal adhesion<br>GO:0005929 (located in) cilium<br>GO:0009986 (located in) cell surface<br>GO:0010008 (located in) endosome membrane<br>GO:0016020 (located in) membrane<br>GO:0016323 (located in) basolateral plasma membrane |

| | | |
|---|---|---|
| | | GO:0016324 (located in) apical plasma membrane<br>GO:0016328 (located in) lateral plasma membrane<br>GO:0019898 (located in) extrinsic component of membrane<br>GO:0030659 (located in) cytoplasmic vesicle membrane<br>GO:0031232 (located in) extrinsic component of external side of plasma membrane<br>GO:0031313 (located in) extrinsic component of endosome membrane<br>GO:0031410 (located in) cytoplasmic vesicle<br>GO:0031514 (located in) motile cilium<br>GO:0031901 (located in) early endosome membrane<br>GO:0031982 (located in) vesicle<br>GO:0042383 (located in) sarcolemma<br>GO:0042995 (located in) cell projection<br>GO:0070062 (located in) extracellular exosome<br>GO:0062023 (part of) collagen-containing extracellular matrix |
| PB0201 (TGF-B3) | 413 | GO:0005114 (enables) type II transforming growth factor beta receptor binding<br>GO:0005125 (enables) cytokine activity<br>GO:0005160 (enables) transforming growth factor beta receptor binding<br>GO:0005515 (enables) protein binding<br>GO:0008083 (enables) growth factor activity<br>GO:0034713 (enables) type I transforming growth factor beta receptor binding<br>GO:0034714 (enables) type III transforming growth factor beta receptor binding<br>GO:0042802 (enables) identical protein binding<br>GO:0044877 (enables) protein-containing complex binding<br>GO:0050431 (enables) transforming growth factor beta binding<br>GO:0001666 (involved in) response to hypoxia<br>GO:0001701 (involved in) in utero embryonic development<br>GO:0007179 (involved in) transforming growth factor beta receptor signaling pathway<br>GO:0007435 (involved in) salivary gland morphogenesis<br>GO:0007565 (involved in) female pregnancy<br>GO:0007568 (involved in) aging<br>GO:0008284 (involved in) positive regulation of cell population proliferation<br>GO:0008285 (involved in) negative regulation of cell population proliferation<br>GO:0010718 (involved in) positive regulation of epithelial to mesenchymal transition<br>GO:0010862 (involved in) positive regulation of pathway-restricted SMAD protein phosphorylation<br>GO:0030501 (involved in) positive regulation of bone mineralization<br>GO:0030509 (involved in) BMP signaling pathway<br>GO:0030512 (involved in) negative regulation of transforming growth factor beta receptor signaling pathway<br>GO:0030879 (involved in) mammary gland development<br>GO:0032570 (involved in) response to progesterone<br>GO:0032967 (involved in) positive regulation of collagen biosynthetic process<br>GO:0034616 (involved in) response to laminar fluid shear stress<br>GO:0042060 (involved in) wound healing<br>GO:0042476 (involved in) odontogenesis<br>GO:0042704 (involved in) uterine wall breakdown<br>GO:0043065 (involved in) positive regulation of apoptotic process<br>GO:0043410 (involved in) positive regulation of MAPK cascade<br>GO:0043524 (involved in) negative regulation of neuron apoptotic process<br>GO:0043627 (involved in) response to estrogen<br>GO:0043932 (involved in) ossification involved in bone remodeling<br>GO:0045216 (involved in) cell-cell junction organization<br>GO:0045893 (involved in) positive regulation of transcription, DNA-templated<br>GO:0045944 (involved in) positive regulation of transcription by RNA polymerase II<br>GO:0048286 (involved in) lung alveolus development<br>GO:0048565 (involved in) digestive tract development<br>GO:0048702 (involved in) embryonic neurocranium morphogenesis<br>GO:0048839 (involved in) inner ear development<br>GO:0050714 (involved in) positive regulation of protein secretion<br>GO:0051491 (involved in) positive regulation of filopodium assembly<br>GO:0051496 (involved in) positive regulation of stress fiber assembly<br>GO:0051781 (involved in) positive regulation of cell division |

| | | |
|---|---|---|
| | | GO:0060325 (involved in) face morphogenesis<br>GO:0060364 (involved in) frontal suture morphogenesis<br>GO:0060391 (involved in) positive regulation of SMAD protein signal transduction<br>GO:0060395 (involved in) SMAD protein signal transduction<br>GO:0062009 (involved in) secondary palate development<br>GO:0070483 (involved in) detection of hypoxia<br>GO:1904706 (involved in) negative regulation of vascular associated smooth muscle cell proliferation<br>GO:1905075 (involved in) positive regulation of tight junction disassembly<br>GO:0005615 (is active in) extracellular space<br>GO:0005576 (located in) extracellular region<br>GO:0005634 (located in) nucleus<br>GO:0005737 (located in) cytoplasm<br>GO:0005886 (located in) plasma membrane<br>GO:0009986 (located in) cell surface<br>GO:0030141 (located in) secretory granule<br>GO:0030315 (located in) T-tubule<br>GO:0031093 (located in) platelet alpha granule lumen<br>GO:0043025 (located in) neuronal cell body<br>GO:0043231 (located in) intracellular membrane-bounded organelle<br>GO:0062023 (located in) collagen-containing extracellular matrix |
| PB0006 (Osteonectin, SPARC) | 303 | GO:0005201 (enables) extracellular matrix structural constituent<br>GO:0005509 (enables) calcium ion binding<br>GO:0005515 (enables) protein binding<br>GO:0005518 (enables) collagen binding<br>GO:0046872 (enables) metal ion binding<br>GO:0050840 (enables) extracellular matrix binding<br>GO:0001937 (involved in) negative regulation of endothelial cell proliferation<br>GO:0010595 (involved in) positive regulation of endothelial cell migration<br>GO:0016525 (involved in) negative regulation of angiogenesis<br>GO:0022604 (involved in) regulation of cell morphogenesis<br>GO:0048856 (involved in) anatomical structure development<br>GO:0005576 (located in) extracellular region<br>GO:0005604 (located in) basement membrane<br>GO:0005615 (located in) extracellular space<br>GO:0005737 (located in) cytoplasm<br>GO:0005886 (located in) plasma membrane<br>GO:0009986 (located in) cell surface<br>GO:0016363 (located in) nuclear matrix<br>GO:0031091 (located in) platelet alpha granule<br>GO:0031092 (located in) platelet alpha granule membrane<br>GO:0031093 (located in) platelet alpha granule lumen<br>GO:0043231 (located in) intracellular membrane-bounded organelle<br>GO:0062023 (located in) collagen-containing extracellular matrix<br>GO:0071682 (located in) endocytic vesicle lumen |

* *Not found in D. rerio*

**Table S12.**
Protein functional domains of the selected proteins

| Protein (Name) | Length | Functional Domains |
|---|---|---|
| PB0249 (BSP)* | 317 | PF05432 (BSP_II) |
| PB0102 (Osteopontin) | 344 | PF00865 (Osteopontin) |
| PB0085 (Fetuin B) | 507 | PF00031 (Cystatin) |
| PB0001 (Fetuin A, AHSG) | 418 | PF00031 (Cystatin) |
| PB0195 (MATN3) | 489 | PF00092 (VWA), PF10393 (Matrilin_ccoil) |
| PB0252 (RSPO1) | 272 | PF00090 (TSP_1)<br>PF15913 (Furin-like_2) |
| PB0092 (ALP) | 564 | PF00245 (Alk_phosphatase) |
| PB0059 (Annexin A1) | 346 | PF00191 (Annexin) |
| PB0201 (TGF-B3) | 413 | PF00019 (TGF_beta), PF00688 (TGFb_propeptide) |
| PB0006 (Osteonectin, SPARC) | 303 | PF00050 (Kazal_1), PF09289 (FOLN), PF10591 (SPARC_Ca_bdg) |

* *Not found in D. rerio*

**Table S13.**
Bone ECM proteins used in mechanical stimulation experiments. Gene ontology and domain annotations of these proteins are available in tables S11 and S12.

| Protein (Name) | Function | Ref. |
|---|---|---|
| PB0249 (BSP) * | Glycoprotein; a molecular marker of mature osteoblasts and osteocytes; plays a role in matrix mineralization; high affinity for calcium; enhances osteoblast differentiation. | (7) |
| PB0102 (Osteopontin) | Glycoprotein; a molecular marker of mature osteoblasts; interact with integrin; involved in cell adhesion, migration and proliferation; suppress osteoblast responses to mechanical stress. | (8,9) |
| PB0085 (Fetuin B) | Glycoprotein; potentially increases serum level with osteoporosis | (10) |
| PB0001 (Fetuin A, AHSG) | Glycoprotein; inhibit ectopic mineralization; low concentration in postmenopausal women with osteoporosis | (11,12) |
| PB0195 (MATN3) | A member of the matrilin family; present in cartilage; regulate ECM degradation; osteoarthritis prevention | (13) |
| PB0252 (RSPO1) | R-spondin family protein; promote osteoblast differentiation via Wnt signaling pathway; inhibit osteoblast apoptosis via activating the Wnt/beta-catenin signaling pathway; upregulated by mechanical stimulation. | (14–16) |
| PB0092 (ALP) | Glycoprotein; a molecular marker of mature osteoblasts; an important enzyme in the process of biomineralization | (17) |
| PB0059 (Annexin A1) | A member of the annexin superfamily; inhibit osteoclast differentiation | (1,18) |
| PB0201 (TGF-β3) | A member of the TGF β superfamily; mechanical loading stimulates the release of TGF-β activity from osteoblasts. | (1,19,20) |
| PB0006 (Osteonectin, SPARC) | Glycoprotein; it binds to collagen type I and hydroxyapatite in bone ECM. | (1,20,21) |

* *Not found in Danio rerio*

**Table S14.**
Expected synonymous and non-synonymous mutations per site.

| 2nd -> | | T | | | | C | | | | A | | | | G | | | |
|---|---|---|---|---|---|---|---|---|---|---|---|---|---|---|---|---|---|
| **1st** | **3rd** | **Codon** | **AA** | **N** | **S** | **Codon** | **AA** | **N** | **S** | **Codon** | **AA** | **N** | **S** | **Codon** | **AA** | **N** | **S** |
| **T** | **T** | TTT | F | 2.67 | 0.33 | TCT | S | 2.00 | 1.00 | TAT | Y | 2.67 | 0.33 | TGT | C | 2.67 | 0.33 |
| | **C** | TTC | F | 2.67 | 0.33 | TCC | S | 2.00 | 1.00 | TAC | Y | 2.67 | 0.33 | TGC | C | 2.67 | 0.33 |
| | **A** | TTA | L | 2.33 | 0.67 | TCA | S | 2.00 | 1.00 | TAA | * | . | . | TGA | * | . | . |
| | **G** | TTG | L | 2.33 | 0.67 | TCG | S | 2.00 | 1.00 | TAG | * | . | . | TGG | W | 3.00 | 0.00 |
| **C** | **T** | CTT | L | 2.00 | 1.00 | CCT | P | 2.00 | 1.00 | CAT | H | 2.67 | 0.33 | CGT | R | 2.00 | 1.00 |
| | **C** | CTC | L | 2.00 | 1.00 | CCC | P | 2.00 | 1.00 | CAC | H | 2.67 | 0.33 | CGC | R | 2.00 | 1.00 |
| | **A** | CTA | L | 1.67 | 1.33 | CCA | P | 2.00 | 1.00 | CAA | Q | 2.67 | 0.33 | CGA | R | 1.67 | 1.33 |
| | **G** | CTG | L | 1.67 | 1.33 | CCG | P | 2.00 | 1.00 | CAG | Q | 2.67 | 0.33 | CGG | R | 1.67 | 1.33 |
| **A** | **T** | ATT | I | 2.33 | 0.67 | ACT | T | 2.00 | 1.00 | AAT | N | 2.67 | 0.33 | AGT | S | 2.67 | 0.33 |
| | **C** | ATC | I | 2.33 | 0.67 | ACC | T | 2.00 | 1.00 | AAC | N | 2.67 | 0.33 | AGC | S | 2.67 | 0.33 |
| | **A** | ATA | I | 2.33 | 0.67 | ACA | T | 2.00 | 1.00 | AAA | K | 2.67 | 0.33 | AGA | R | 2.67 | 0.33 |
| | **G** | ATG | M | 3.00 | 0.00 | ACG | T | 2.00 | 1.00 | AAG | K | 2.67 | 0.33 | AGG | R | 2.67 | 0.33 |
| **G** | **T** | GTT | V | 2.00 | 1.00 | GCT | A | 2.00 | 1.00 | GAT | D | 2.67 | 0.33 | GGT | G | 2.00 | 1.00 |
| | **C** | GTC | V | 2.00 | 1.00 | GCC | A | 2.00 | 1.00 | GAC | D | 2.67 | 0.33 | GGC | G | 2.00 | 1.00 |
| | **A** | GTA | V | 2.00 | 1.00 | GCA | A | 2.00 | 1.00 | GAA | E | 2.67 | 0.33 | GGA | G | 2.00 | 1.00 |
| | **G** | GTG | V | 2.00 | 1.00 | GCG | A | 2.00 | 1.00 | GAG | E | 2.67 | 0.33 | GGG | G | 2.00 | 1.00 |
| | | | | | | | | | | | | | | --- | 0 | 3.00 | 0.00 |

1st: nucleotide in the first position of the codon; 2nd: nucleotide in the second position of the codon; 3rd: nucleotide in the third position of the codon; AA: amino acid; N: sum of non-synonymous mutations expected per site; S: sum of synonymous mutations expected per site; -: indel; *: stop codon.
